# Functional analysis of phospholipase Dδ family in tobacco pollen tubes

**DOI:** 10.1101/807719

**Authors:** Přemysl Pejchar, Juraj Sekereš, Ondřej Novotný, Viktor Žárský, Martin Potocký

## Abstract

Phosphatidic acid (PA), important signalling and metabolic phospholipid, is predominantly localized in the subapical plasma membrane (PM) of growing pollen tubes. PA can be produced from structural phospholipids by phospholipase D (PLD) but the isoforms responsible for production of plasma membrane PA were not identified yet and their functional roles remain unknown. Following genome-wide bioinformatic analysis of PLD family in tobacco, we focused on the pollen-overrepresented PLDδ class. Combining live-cell imaging, gene overexpression or knock-down, lipid-binding and structural bioinformatics, we characterized 5 NtPLDδ isoforms. Distinct PLDδ isoforms preferentially localize to the cytoplasm or subapical PM. Using fluorescence recovery after photobleaching, domain deletion and swapping analyses we show that membrane-bound PLDδs are tightly bound to PM, primarily via the central catalytic domain. Knock-down, overexpression and *in vivo* PA level analyses revealed isofom PLDδ3 as the most important member of the PLDδ subfamily active in pollen tubes. PA promotes binding of PLDδ3 to the PM, thus creating a positive feedback loop, where PA accumulation leads to the formation of massive PM invaginations. Tightly controlled production of PA generated by PLDδ3 at the PM is important for maintaining the balance between various membrane trafficking processes, that are crucial for plant cell tip growth.

## Introduction

Polar cell growth is one of the most fundamental processes in plant development. It determines cellular morphogenesis and ultimately defines plant phenotype (Cole & Fowler, 2006; Sekereš *et al.*, 2015). Because it is challenging to study the control of cell polarity in tissues or organs composed of multiple interacting cell types, readily accessible, extremely polarized and fast-growing cells could be beneficially utilized as cell morphogenesis model systems. Pollen tubes, formed as an extremely elongated outgrowth upon germination of a pollen grain, have an indispensable function in the fertilization of higher plants. Pollen tube tip growth is a widely used model perfectly suited to study cellular processes that underlie polarized cell expansion (Grebnev *et al.*, 2017). Furthermore, other tip-growing cells like root hairs, moss protonemal cells, fungal hyphae and even neuronal axons share many conserved molecular mechanisms for establishing and maintaining cell polarity with pollen tubes, despite their evolutionary distance (Lee *et al.*, 2008; Ischebeck *et al.*, 2010; Žárský & Potocký, 2010; Vaškovičová *et al.*, 2013).

The cell polarity is primarily manifested by the asymmetric distribution of proteins and lipids in the specialized domains of the plasma membrane (PM), an asymmetric proteolipidic structure containing roughly 50–100 molecules of lipids per protein (Furt *et al.*, 2011). Around 3000 proteins is found in the PM proteome (Cao *et al.*, 2016), and its lipidome contains thousands of molecular species (van Meer *et al.*, 2008). PM polar domains are formed by balanced exocytosis, endocytosis, endomembrane trafficking, lateral mobility and by distinctly localized peripheral phospholipid-modifying enzymes (Žárský & Potocký, 2009; Orlando & Guo, 2009; Gronnier *et al.*, 2018). Several anionic phospholipids, namely polyphosphoinositides phosphatidylinositol 4-phosphate (PI4P) and phosphatidylinositol 4,5-bisphosphate (PIP_2_), together with phosphatidylserine (PS) and phosphatidic acid (PA) define PM identity by maintaining electronegative electrostatic properties, by modulating PM biophysical properties and via specific targeting of effector proteins to the cytoplasmic leaflet of PM (Heilmann & Ischebeck, 2015; Noack & Jaillais, 2017). In tip-growing cells, laterally separated PM domains are distinguishable by different enrichment of specific phospholipids and/or proteins (Potocký *et al.*, 2014; Platre *et al.*, 2018). Notably, phospholipid-modifying enzymes, peripherally attached to PM, are crucial for this patterning. PIP_2_ localizes near the tip of growing pollen tubes and this narrow localization is maintained by a combination of two counteracting enzyme activities: near-apically localized phosphatidylinositol 4-phosphate 5-kinases (PIP5Ks), which produce PIP_2_ along the broad apical region and subapically-located phospholipases C (PLCs), that focus PIP_2_ localization to the tip by cleaving it into DAG and inositol trisphosphate (Helling *et al.*, 2006; Ischebeck *et al.*, 2008, 2011).

Phosphatidic acid (PA) is the simplest phospholipid with just a phosphate headgroup. Despite its simple structure, PA is important in many processes of all eukaryotic cells. It serves as a key intermediate in the phospholipid biosynthetic pathways and plays an important function in vesicular traffic and cytoskeletal dynamics. PM levels of PA were reported in range of 4-16% (Furt *et al.*, 2011). We previously described that PLD-produced PA is required for polar expansion of pollen tubes (Potocký *et al.*, 2003) and that PA depletion decreases the number of secretory vesicles in the pollen tube tip (Pleskot *et al.*, 2012). Two major, distinct signalling pathways contribute to the production of PA at the PM (Pleskot *et al.*, 2014). One is a direct cleavage of structural phospholipids by phospholipase D (PLD), and another is a sequence of reactions catalyzed by phospholipase C (PLC) and diacylglycerol kinase (DGK).

In comparison to yeast and animal genomes, the PLD family is expanded in plants with 12 genes in Arabidopsis and even more in other genomes (Eliáš *et al.*, 2002; Pleskot *et al.*, 2012). Canonical plant PLDs can be divided into two major subfamilies: (*i*) PXPH-type PLDs possessing the PHOX homology (PX) and pleckstrin homology (PH) domains that are related to animal/fungal enzymes and (*ii*) plant-specific C2-PLDs characterized by the presence of the N-terminal C2 domain. The plant-specific C2-PLD subfamily members have been further divided into several classes (PLDα, β, γ, ε and δ) based on their phylogeny, exon-intron and domain structures, biochemical properties and substrate preferences (Li *et al.*, 2009; Testerink & Munnik, 2011; Takáč *et al.*, 2019). In Arabidopsis, PLDα1 was shown to participate in many (mostly abiotic) stress responses, including ABA, ethylene and jasmonate signalling pathways, while PLDβ1 isoform has been implicated mainly in responses to biotic stress (Hong *et al.*, 2016; Li & Wang, 2019). PLDγ1 is involved in aluminium stress signalling (Zhao *et al.*, 2011), while PLDε then may play a role in connecting membrane sensing of nitrogen availability to the regulation of growth (Hong *et al.*, 2009). Arabidopsis single PLDδ isoform is highly expressed in most tissues and its transcript is further elevated during drought, salt stress and cold acclimation. Upregulation of PLDδ in transgenic Arabidopsis positively regulates plant tolerance to freezing, oxidative assault, and UV radiation, and genetic analysis implicated PLDδ in response to pathogens (Pinosa *et al.*, 2013; Hong *et al.*, 2016). Interestingly, PLDδ can also be activated by H_2_O_2_ and was shown to physically interact with both microtubular and actin cytoskeleton (Zhang, 2003; Andreeva *et al.*, 2009; Potocký *et al.*, 2012). In PHPX-PLD subfamily, PLDζ1 and PLDζ2 have been shown to participate in auxin-mediated root tropism and phosphate deficiency response (Mancuso *et al.*, 2007; Li & Xue, 2007; Korver *et al.*, 2019).

Despite the long history of plant PLD research and wealth of biochemical and physiological observations, comprehensive data about PLD localization and their mode of action at the PM are still missing. Only Arabidopsis PLDε and δ were shown to be PM-bound (Hong *et al.*, 2009; Pinosa *et al.*, 2013). Similarly, very little isoform-specific data exist for the involvement of PLD in tip growth, despite pharmacological observations (Potocký *et al.*, 2003; Monteiro, 2005; Pleskot *et al.*, 2012). Interestingly, the interaction of PLDβ with actin and production of PA play an important role in pollen tube growth (Pleskot *et al.*, 2010). Here we show that angiosperm pollen tubes have maintained high expression of PLDδ class genes and we present a comprehensive analysis of five PLDδ isoforms in growing tobacco pollen tubes. We demonstrated a distinct localization and membrane dynamics for individual PLDδs and identified sequence features that are important for PM binding of selected PLDδ isoforms. We observed that elevated levels of PA produced by overexpression of the major pollen isoform NtPLDδ3 induced specific morphological phenotypes, indicating a disturbed balance in vesicular trafficking and membrane recycling.

## Results

### Phylogenetic and expression analysis of phospholipases D shows that PLDδ class expanded in angiosperms and is overrepresented in pollen

In order to establish which PLD isoforms are present in tobacco pollen, we first analyzed the phylogeny of PLD gene family in genus *Nicotiana*. We performed homology searches in tobacco genome drafts available at NCBI and <u>solgenomics.net</u>, using Arabidopsis PLDs as input sequences. Additionally, draft genomes of tomato (*Solanum lycopersicum*), *Nicotiana sylvestris, Nicotiana tomentosiformis, Nicotiana benthamiana* and *Solanum tuberosum* were also searched. Following this strategy we identified 30 genes coding for canonical PLDs in allotetraploid tobacco genome that correspond to 15 PLDs in the diploid gene set. The sequences of PLD homoeologous gene pairs originating from parental genomes are nearly identical (>96% nucleotide identity) and previous analyses showed neither differences in gene expression nor the creation of a new function for homoeologous genes in *N. tabacum* parental subgenomes (Sierro *et al.*, 2013; Edwards *et al.*, 2017). For the sake of clarity, in subsequent analyses we considered only one gene per homoeologous pair, i.e. 15 PLD genes that belong to parental subgenome datasets.

To get better insight into the evolution of tobacco PLDs, we performed phylogenetic and expression analyses of PLD family in 5 diverse angiosperm species (tobacco, tomato, Arabidopsis, poplar, grapevine and rice), for which extensive sporophyte and gametophyte transcriptomic data are available. Fig. 1a and Table 1 show that all five previously described PLD classes, i.e. α, β/γ, δ, ε belonging to C2-PLD subfamily and ζ class from PXPH-PLD subfamily, (see Eliáš *et al.* (2002), Qin & Wang (2002) and Pleskot *et al.* (2012) for more details) are present in tobacco genome. Despite many independent gene duplications or losses detectable in most of the species, total gene number in individual PLD subclasses is quite stable, although the distribution of isoforms within certain classes (in particular α and δ) differs (Fig. 1a). One notable exception is Arabidopsis, where on one hand PLDδ (typically quite expanded and represented by 3-5 members in other species) was reduced to a single gene but on the other hand PLDγ class is present only in Arabidopsis (and other Brassicaceae, data not shown).

**Table 1.** Gene repertoire for various PLD classes in selected angiosperm genomes and transcript presence in pollen

| | PLDα | | PLDε | | PLDβ | | PLDγ | | PLDδ | | PLDζ | |
| --- | --- | --- | --- | --- | --- | --- | --- | --- | --- | --- | --- | --- |
|  | *genes* | *pollen* | *genes* | *pollen* | *genes* | *pollen* | *genes* | *pollen* | *genes* | *pollen* | *genes* | *pollen* |
| *Nicotiana tabacum*<sup>1</sup> | 4 | 1 | 1 | 0 | 3 | 1 | - | - | 5 | 5 | 2 | 1 |
| *Solanum lycopersicum* | 4 | 1 | 1 | 0 | 3 | 1 | - | - | 5 | 4 | 2 | 1 |
| *Arabidopsis thaliana* | 3 | 1 | 1 | 1 | 2 | 2 | 3 | 2 | 1 | 1 | 2 | 1 |
| *Populus trichocarpa* | 4 | 3 | 1 | 0 | 3 | 2 | - | - | 5 | 5 | 2 | 2 |
| *Vitis vinifera* | 4 | 2 | 1 | 1 | 2 | 2 | - | - | 3 | 3 | 2 | 1 |
| *Oryza sativa* | 7 | 1 | 1 | 0 | 2 | 2 | - | - | 4 | 4 | 2 | 2 |

*genes* - total number of genes belonging to individual isoforms. *pollen* - number of genes expressed in pollen (detected in RNA-Seq or Affymetrix assay). <sup>1</sup>for the sake of clarity, only data from one diploid subgenome are shown.

**Fig. 1.**
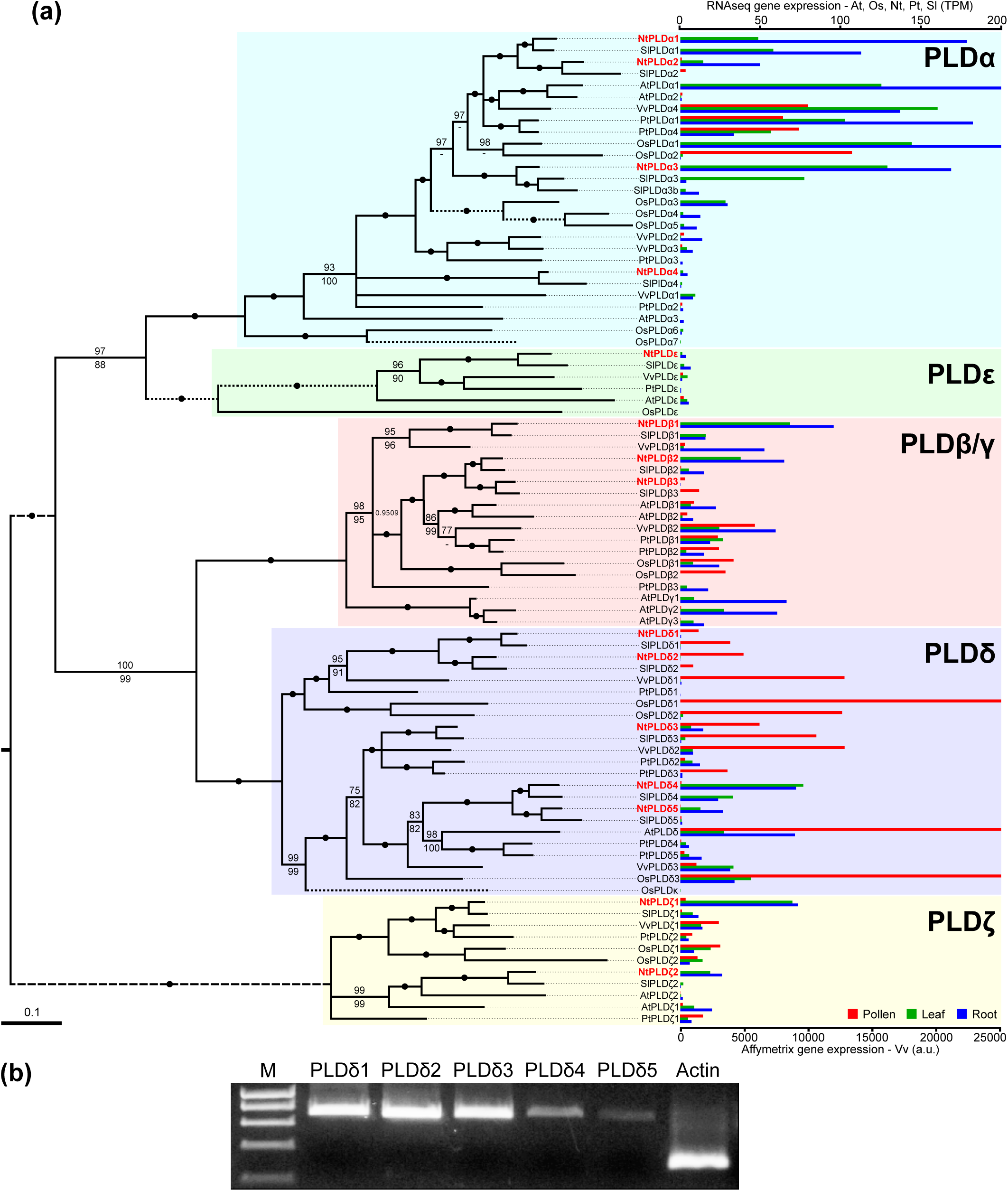
PLDδ subfamily has expanded in most plant species and is expressed preferentially in pollen. **(a)** Phylogenetic and transcriptomic analysis of canonical plant PLDs from six diverse angiosperm species. On the left, phylogenetic tree represents protein maximum likelihood (ML) phylogeny rooted with human PLD2 (not shown). Numbers at nodes correspond to the aLRT test with the SH-like (Shimodaira–Hasegawa-like) support from ML (top) and posterior probabilities from Bayesian analysis (bottom). Nodes with 100% support from both algorithms are marked by black dots. Missing values indicate support below 50%. Branches were collapsed if the inferred topology was not supported by both methods. Scale bar indicates the rates of substitutions/site. For the sake of clarity, some branches were shortened to ½ (dotted line) or ¼ (dashed line) of their original length. On the right, relative gene expression levels from pollen and two sporophytic tissues are shown, using publicly available RNA-seq or Affymetrix microarray data. **(b)** Semiquantitative RT-PCR analysis confirming that all five *NtPLDδ* genes are expressed in pollen tubes. Species abbreviations: At, *Arabidopsis thaliana*; Nt, *Nicotiana tabacum*; Os, *Oryza sativa*; Pt, *Populus trichocarpa*; Sl, *Solanum lycopersicum*; Vv, *Vitis vinifera*. PLD, phospholipase D.

We then analyzed PLD expression in pollen by remapping tobacco pollen raw RNA-seq data (Conze *et al.*, 2017) on reference genome sequence and compared it to the publicly available pollen, leaf and root RNA-seq data from other species (Fig. 1a and Table 1). This clearly demonstrated that distinct PLD classes have very conserved tissue expression in angiosperms and that genes from PLDδ class (i.e. all 5 tobacco isoforms) show high pollen expression values. No other PLD class exhibited the same pattern. Close-up inspection of the data revealed the presence of two evolutionary conserved subgroups within PLDδ class: the first one (that includes *NtPLDδ1* and *NtPLDδ2*) is nearly pollen exclusive, while the second one (that includes *NtPLDδ3, NtPLDδ4* and *NtPLDδ5*) has more diverse expression profile. To experimentally confirm the expression of *NtPLDδ* genes in pollen we performed RT–PCR showing that transcripts of all five *NtPLDδ* genes were indeed detected in pollen tubes with the high level for *NtPLDδ1-3* and low for *NtPLDδ4* and *NtPLDδ5*, nicely corroborating RNA-seq data (Fig. 1b).

Collectively, our genome-wide phylogeny and expression analysis strongly suggested that PLDδ class represents the majority of PLD isoforms expressed in the male gametophyte throughout angiosperms evolution.

### Distinct PLDδ members show different localization patterns and membrane dynamics

Having established PLDδ as the dominant PLD class in tobacco male gametophyte, we focused on the localization of individual PLDδ isoforms in growing pollen tubes. We fused NtPLDδs and AtPLDδ with YFP under the control of pollen-specific *Lat52* promoter and monitored the localization of YFP:PLDδs using spinning disk confocal microscopy and measured pollen tube mean growth rates in transiently transformed pollen tubes. Individual isoforms of NtPLDδs and AtPLDδ were found to have distinct fluorescent signal, that reflected their phylogenetic relations. YFP:NtPLDδ1 and YFP:NtPLDδ2 showed cytoplasmic localization devoid of any distinguishable subcellular features. Conversely, YFP:NtPLDδ3-5 and YFP:AtPLDδ decorated subapical PM starting ∼10-15 µm behind the tip. Interestingly the apical PM signal was never observed at the extreme apex during cell elongation (Fig. 2). YFP:NtPLDδ3 showed only faint PM signal exclusively in subapical zone, while PM signals of YFP:NtPLDδ4-5 and YFP:AtPLDδ were more pronounced and extended also further back to the pollen tube shank (Fig. 2). We confirmed the distinct PM/cytoplasmic localization of YFP:NtPLDδ isoforms by co-staining with PM/endocytic dye FM4-64 and by the imaging of PLDδ C-terminal YFP fusions, which also ruled out the possible effect of YFP position (Fig. S1).

**Fig. 2.**
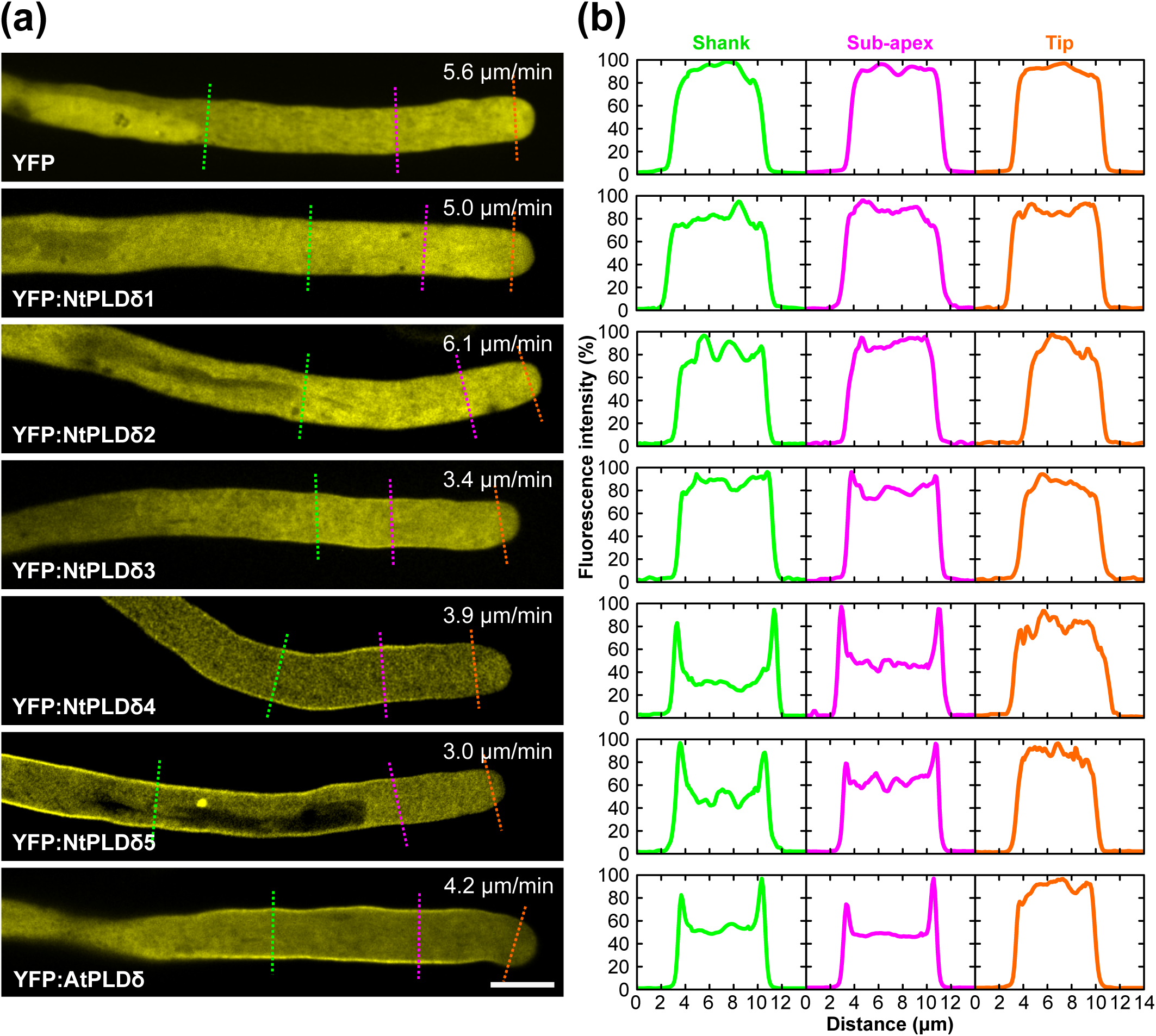
Tobacco and Arabidopsis PLDδ isoforms show distinct localization patterns in tobacco pollen tubes. **(a)** Localization of YFP-tagged NtPLDδ1-5 and AtPLDδ in actively growing pollen tubes (PTs). Pollen was transiently transformed with 1 μg of DNA and analyzed 8 to 10 h after transformation. Numbers inserted in figures indicate PT growth rates. Bar, 10 μm. **(b)** Relative fluorescence intensity across the PT tip (orange lines), across the PT subapex (purple lines) or across the PT shank (green lines) for pollen tubes shown in (a). Fluorescence was measured using the line scan tool in ImageJ across 14 μm-long regions indicated in (a) with line width set to 5 pixels. PLD, phospholipase D.

The observed differences in PM localization for YFP:NtPLDδ3-5 and YFP:AtPLDδ led us to assess their dynamic behaviour at the PM using Fluorescence Recovery After Photobleaching (FRAP) assay. We bleached small subapical region of PM in pollen tubes expressing PM-bound NtPLDδs or AtPLDδ and followed the recovery using confocal microscopy. Given that membrane-bound PLDδs are peripheral proteins with substantial cytoplasmic fraction, we expected a fast fluorescence recovery, however the kymograph projections indicated surprisingly slow dynamics (Fig. 3a). Indeed, the quantitative analysis of FRAP curves confirmed this and revealed that a large proportion of membrane PLDδs resides at the PM, reaching immobile fraction up to ∼70% for NtPLDδ5 and AtPLDδ. To estimate the relative involvement of slow lateral PM diffusion and fast PM/cytoplasm exchange of individual PLDδ isoforms, we performed a simple analysis by fitting the FRAP curves with either single or double exponential (Fig. 3bc, Fig. S2). We validated this approach using cells expressing laterally diffusing, PM-anchored lipidated YFP (MAP-YFP), or low-affinity lipid-binding domain (Cys1-YFP) that predominantly shuttles quickly between PM and cytoplasm. In both cases, single exponential fitted the data well, showing typical values for fast (Cys1-YFP) or slow (MAP-YFP) movement. Analysis of PLDδ curves suggested that lateral PM diffusion is major component (∼60-100%) of PLDδ dynamics, particularly for YFP:NtPLDδ5 and YFP:AtPLDδ. Altogether, these experiments showed that all membrane-bound PLDδ isoforms, and NtPLDδ5 and YFP:AtPLDδ in particular, are remarkably strongly attached to PM, with very low shuttling to/from cytoplasm.

**Fig. 3.**
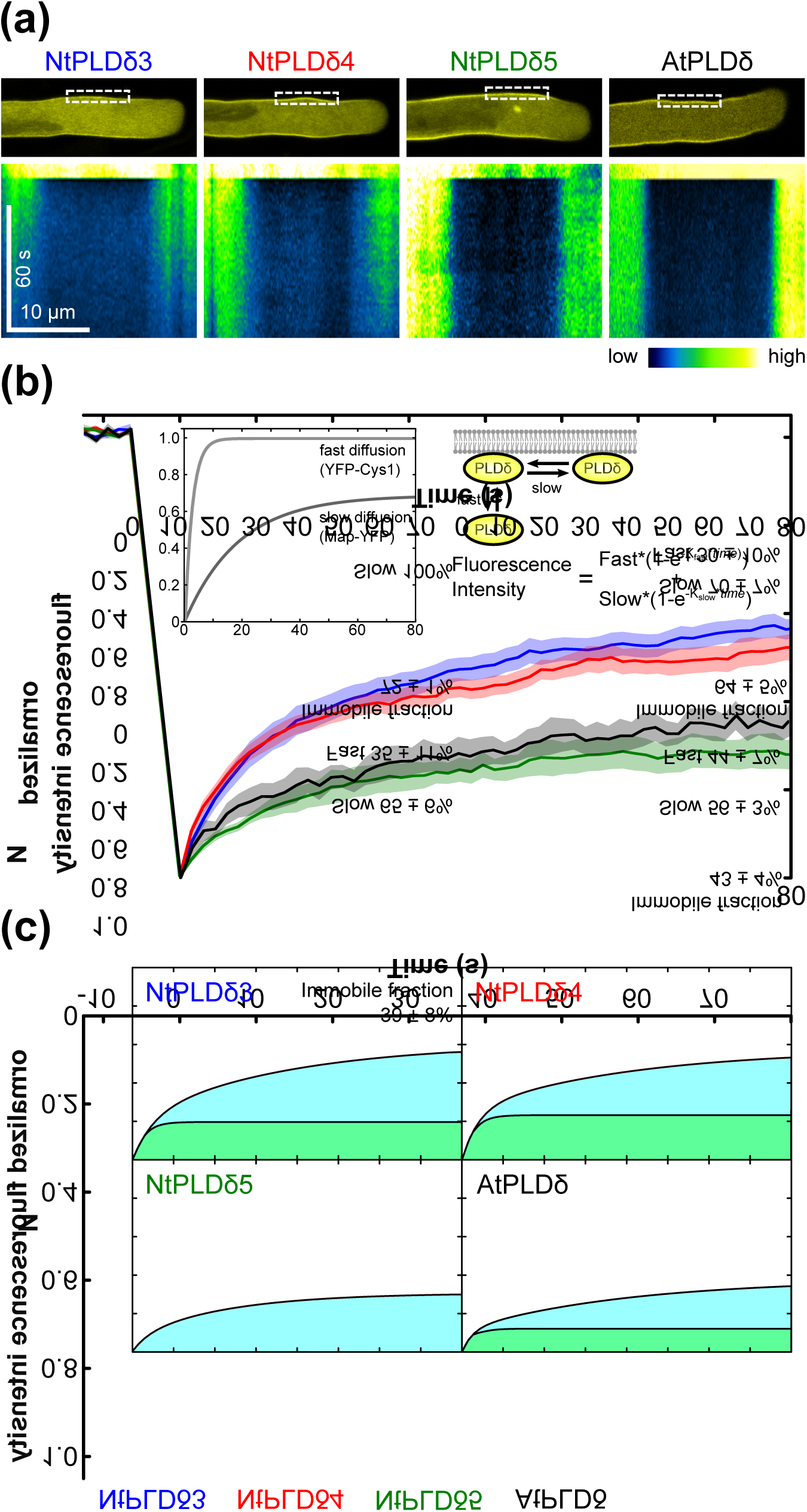
FRAP analyses show distinct cellular dynamics for membrane-bound members of PLDδ family. **(a)** Representative confocal images showing the bleached ROI and kymographs showing the fluorescence recovery at the plasma membrane (PM). (**b**) Quantitative analysis of YFP:NtPLDδ3-5 and YFP:AtPLDδ FRAP at pollen tube PM. Each curve shows mean normalized fluorescence intensity (n = 10-18 in two separate experiments), SEM is indicated by the shaded area. Inset picture shows the schematic representation of major factors contributing to PM fluorescence recovery and formula used for fitting the experimental data. Inset chart shows FRAP curves of peripheral membrane proteins showing only fast PM-cytosol exchange or slow lateral movement along PM, respectively. See Fig. S2 for more details. **(c)** Analysis of fitted FRAP curves showing immobile fractions at the PM and relative contributions of fast and slow components to PLDδ mobile fraction. PLD, phospholipase D.

### N-terminal and central catalytic domain are required for correct intracellular targeting of NtPLDδ isoforms

Having observed the distinct localization of NtPLDδ1-2 vs NtPLDδ3-5, we investigated in detail the molecular determinants responsible for PM binding of certain NtPLDδs. We chose the most PM-localized isoform NtPLDδ4 as a representative of PM NtPLDδs and NtPLDδ2 as typically cytoplasmic protein. First, we prepared a set of YFP:NtPLDδ4 truncations, NtPLDδ4-N (aa 1-180), NtPLDδ4-C (aa 751-869), central/catalytic domain NtPLDδ4-ΔNΔC (aa 181-750), NtPLDδ4-ΔC (aa 1-750) and NtPLDδ4-ΔN (aa 181-869) and monitored their localizations on PM in transiently transformed tobacco pollen tubes (Fig. 4a and Fig. S3). None of these simple truncations were able to decorate PM (Fig. 4b,e) suggesting that both N-terminal and C-terminal parts are necessary for correct PM localization of NtPLDδ4. We next combined N-terminal and C-terminal part together (NtPLDδ4-NC) but again only cytosolic fluorescent signal was observed. Taking into account possible misfolding due to missing domains of the NtPLDδ4 molecule, we substituted these domains with those from cytoplasmic NtPLDδ2 to maintain overall molecule structure. We prepared all possible combinations of chimeric constructs fused to YFP and followed their localization. Surprisingly, only YFP:NtPLDδ442 showed clear signal at PM, suggesting that the catalytic domain is required, but not sufficient, for PM localization of NtPLDδ4 (Fig. 4c,e) and while the C-terminal domain seems not to be involved in direct interaction with PM.

**Fig. 4.**
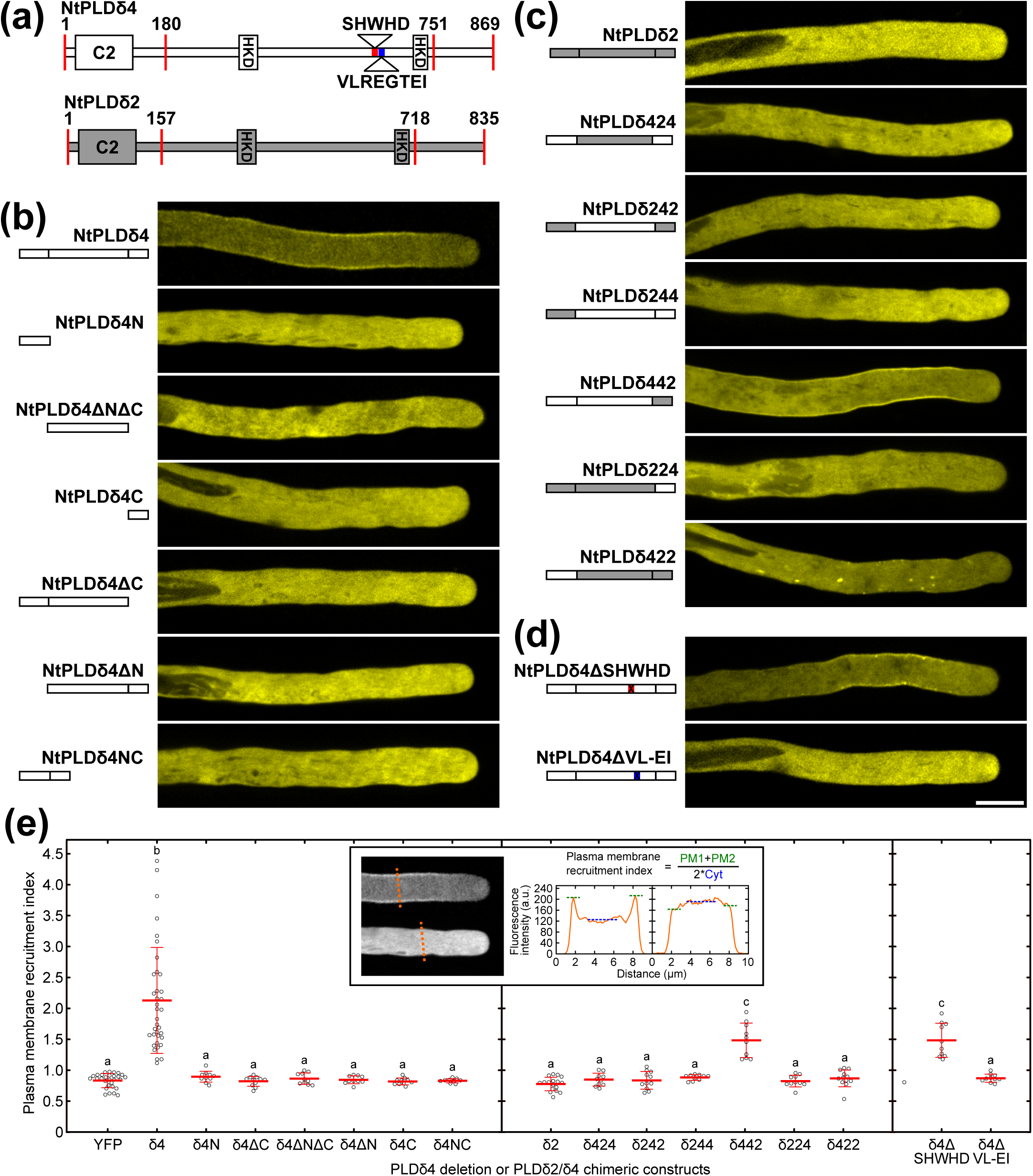
Catalytic domain is required, but not sufficient, for plasma membrane localization of NtPLDδ4. **(a)** Domain organization and subfamily-specific features for NtPLDδ4 and NtPLDδ2 sequences (see Fig. S3 for more details). **(b-d)** The localization analysis of N-terminally YFP-tagged NtPLDδ4 deletion fragments and NtPLDδ2/δ4 chimeric proteins. Schematic representation of the deletion or chimeric construct and typical localization in growing pollen tube is always shown. Bar, 10 μm. **(e)** Quantitative analysis of membrane recruitment of the deletion and chimeric constructs shown in (b-d). Inset depicts the measurement and computation of membrane recruitment index, that was used as the quantitative proxy for membrane association. For each construct, 10-31 cells were imaged in 2-3 independent experiments. Individual data points and mean ± SD are shown, different letters indicate significant differences between samples (P < 0.001). Cyt, cytoplasm; PLD, phospholipase D; PM, plasma membrane.

To further narrow down the determinants of PM-binding within the central catalytic domain, we found (based on the PLDδ multiple alignment) two short motifs that are missing in cytosol-localized NtPLDδ1-2 but present in PM-localized NtPLDδ3-5/AtPLDδ (motif SHWHD in NtPLDδ4) or in NtPLDδ4-5/AtPLDδ only (motif VLREGTEI in NtPLDδ4, Fig. 4a and Fig. S3). We prepared variants of YFP:NtPLDδ4 where these motifs were deleted (NtPLDδ4-ΔSHWHD and NtPLDδ4-ΔVL-EI), and monitored their localization. While the localization and PM-binding of YFP:NtPLDδ4-ΔSHWHD was similar to full-length YFP:NtPLDδ4 (Fig. 4d,e), deletion of the VLREGTEI motif from NtPLDδ4 led to complete loss of the PM localization (Fig. 4d,e).

To get more insight into the complex nature of NtPLDδs interaction with PM, we constructed 3D homology models of selected NtPLDδs to cover all three different types of cellular localization (NtPLDδ2-4, see Fig. 2a). Despite the lack of high-quality structural template, the need for partially *ab initio* modelling, and only an intermediate level of sequence similarity among NtPLDδ2-4 (55-65%), the best models for all three proteins converged to similar tertiary structures with adequate evaluation scores. Modelling suggested that in general, PLDδ tertiary structure consists of tightly packed globular catalytic domain with attached C-terminal domain and somewhat loosely connected N-terminal C2 domain (Fig 5a). The mapping of electrostatic potential revealed that the overall distribution of surface charge is similar for NtPLDδ2-4. However, NtPLDδ4 contains strongly positively charged pocket that surrounds active site (Fig. 5a). Remarkably, short amino acid motif VLREGTEI that was found to be crucial for NtPLDδ4 PM binding (Fig. 4d,e), is localized at NtPLDδ4 molecule surface in close proximity to N-terminal domain where it may be involved in the cooperation of N-terminal and central domains for proper NtPLDδ4 PM targeting (Fig. 4c,e).

**Fig. 5.**
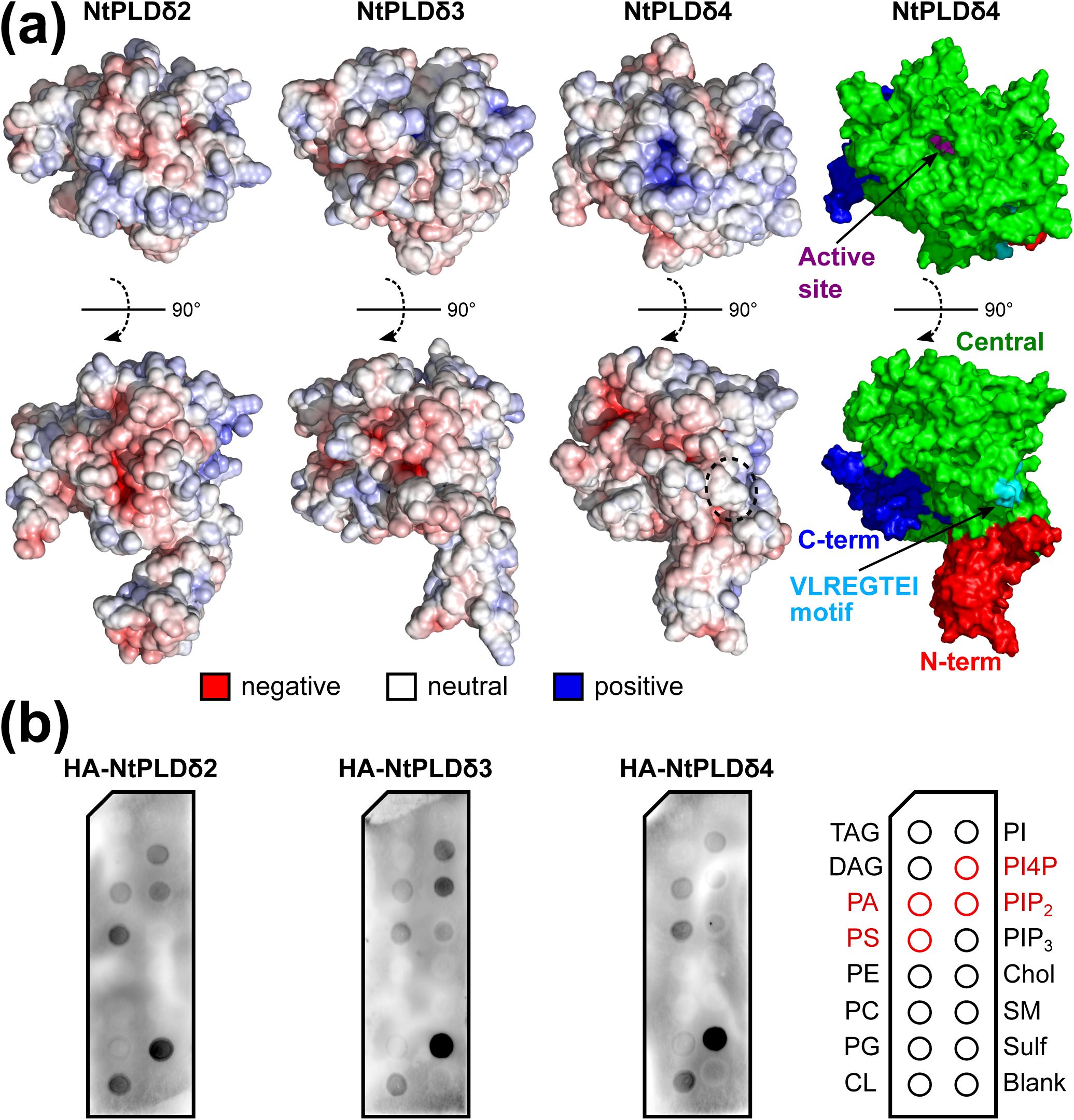
Comparative structural analysis of membrane-binding regions on NtPLDδ isoforms with different membrane-binding properties. **(a)** Electrostatic potentials (left) mapped on the surfaces of the best homology models for NtPLDδ2, NtPLDδ3 and NtPLDδ4 and the schematic depiction (right) of three major domains in NtPLDδ4 homology model (N-terminal C2 domain in red, central catalytic domain in green and C-terminal domain in blue). **(b)** Lipid-binding properties of *in vitro* translated HA-tagged NtPLDδ2, NtPLDδ3 and NtPLDδ4 proteins, as determined using a protein lipid overlay assay. Chol, cholesterol; CL, cardiolipin; DAG, diacylglycerol; PA, phosphatidic acid; PC, phosphatidylcholine; PE, phosphatidylethanolamine; PG, phosphatidylglycerol; PI, phosphatidylinositol; PI4P, phosphatidylinositol 4-phosphate; PIP_2_, phosphatidylinositol 4,5-bisphosphate; PIP_3_, phosphatidylinositol 3,4,5-trisphosphate; PLD, phospholipase D; PS, phosphatidylserine; SM, sphingomyelin; Sulf, sulfatide; TAG, triacylglycerol.

We have previously shown that different membrane lipids localize to distinct domains of pollen tube PM (Potocký *et al.*, 2014), we therefore tested whether the different localization may be due to the ability of NtPLDδs to interact with different spectrum of membrane lipids. We prepared HA-tagged constructs for *in vitro* transcription/translation of NtPLDδ2-4 and used them for lipid-binding assay experiments. In contrast to their different localization, all selected NtPLDδs possess similar ability to bind multiple lipids *in vitro* with strong preference towards negatively charged phospholipids enriched in PM (Fig. 5b). Based on these results, it seems that NtPLDδs are not targeted to PM by specific membrane lipids but possibly another protein interacting partner is needed for proper PM localization.

Taken together, our data show that PM-targeting of selected PLDδ isoforms is a complex process that involves most of the protein and probably integrates several independent protein domains and dynamic steps.

### NtPLDδ3 is the major PLDδ class isoform in tobacco pollen

To get a first insight into the role of NtPLDδ isoforms in pollen tube growth, we studied phenotypes of pollen tubes transiently expressing various levels of YFP-tagged NtPLDδ1-5 or free YFP using confocal microscopy. While overexpression of all NtPLDδ isoforms ultimately resulted in growth retardation, the severity of the response and cellular architecture differed significantly between isoforms (Fig. 4a). Overexpression of YFP:NtPLDδ1-2 did not cause substantial changes in cell morphology and no aberrant intracellular structures were observed. On the other hand, overexpression of PM-bound isoforms YFP:NtPLDδ3-5 led to pronounced alterations of pollen tube structure, where either apical invaginations or accumulation of round intracellular structures were frequently observed (Fig. 6a). This was accompanied by relocalization of YFP:NtPLDδ3-5 to the apical part of cell (Fig. 6a and Fig. S4). Notably, both apical invaginations and intracellular structures were clearly decorated by the fluorescent signal of corresponding isoforms, suggesting that observed intracellular structures had plasma membrane origin. When we quantified the frequency of three major pollen tube tip phenotypes (normal cell, apical invaginations and intracellular membranes), we observed clear difference between YFP:NtPLDδ3 and YFP:NtPLDδ4-5: the overexpression of YFP:NtPLDδ3 resulted in equal occurrence of normal phenotype and apical invaginations but never in formation of the intracellular structures. On the other hand, expression of YFP:NtPLDδ4 and YFP:NtPLDδ5 produced all three phenotypes with the lowest quantities of apical invaginations. Overexpression of each YFP:NtPLDδ isoform caused significant reduction of pollen tube growth compared to overexpressed YFP control with the strongest inhibitory effect detected for YFP:NtPLDδ3 (Fig. 6c). To further confirm the outcomes of the overexpression studies, we employed antisense knock-down strategy which is used to specifically knock-down gene expression in pollen tubes (Potocký *et al.*, 2019). We designed specific antisense oligodeoxynucleotides (ODNs) against all NtPLDδ genes. Addition of liposomes containing NtPLDδ1-3 antisense ODNs into the germination medium significantly retarded pollen tube growth compared to its sense ODNs while we did not observed such effect for NtPLDδ4-5 (Fig. 6d). Also here the strongest effect was observed for NtPLDδ3 with the decrease of pollen tube length to 25% of control. Based on these data it is possible to assume that all NtPLDδ isoforms are involved in pollen tube growth and that NtPLDδ3 is the major functional PLDδ isoform in tobacco pollen.

**Fig. 6.**
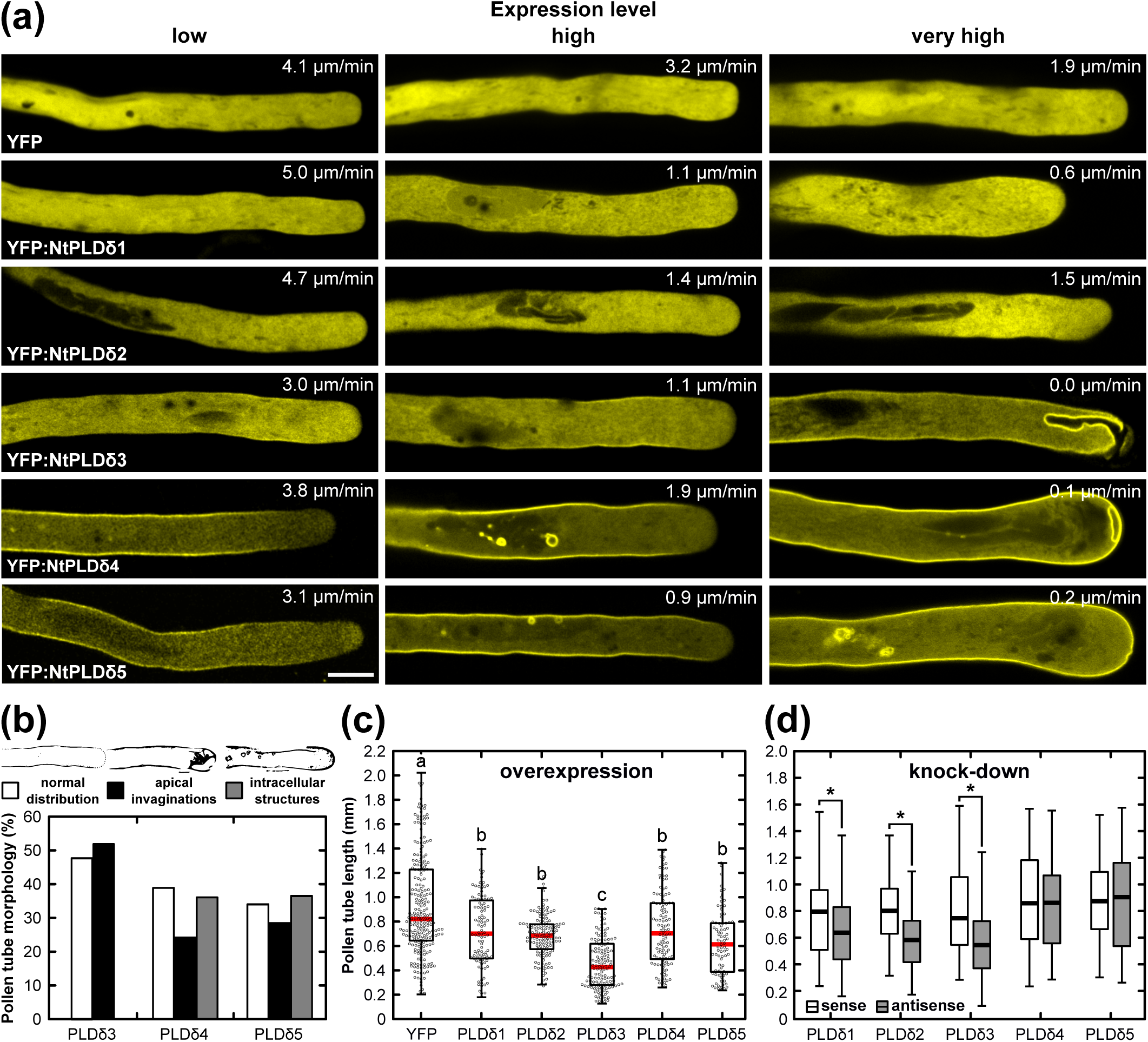
Isoforms of NtPLDδ subfamily show distinct overexpression and knock-down phenotypes in tobacco pollen tubes. **(a)** Typical localization and phenotypes of pollen tubes expressing various levels of free YFP and YFP-tagged NtPLDδ1-5. Pollen was transformed with 0.5 (for low expression level) or 5 μg (for high expression levels) of DNA and analyzed 8 to 10 h after transformation. Numbers inserted in figures indicate pollen tube growth rates. Note that the fluorescence intensities of cells with different expression levels were adjusted to give maximum signal clarity, see Fig. S7 for examples of non-adjusted images. Bar, 10 μm. **(b)** Quantitative analysis of the effect of YFP:NtPLDδ3-5 overexpression on membrane deformations (n > 60, three independent experiments). Cellular phenotypes were divided into three categories indicated by typical examples. **(c)** The effect of NtPLDδ1-5 overexpression on pollen tube growth. Pollen was biolistically transformed with 1 μg of DNA and analyzed 24 h after transformation (n > 80, two independent experiments). Different letters indicate significant differences between samples (P < 0.001). **(d)** The effect of NtPLDδ1-5 knock-down on pollen tube growth. Pollen tubes were cultivated in the presence of 30 μM antisense or sense oligodeoxynucleotides against NtPLDδ1-5 isoforms for 3 h (n > 100, two independent experiments). Asterisk shows significant difference between corresponding antisense-sense pairs (p < 0.01). PLD, phospholipase D.

### PLD isoforms overexpression effect on pollen tubes PA levels at the PM using genetically-encoded PA sensor

We previously showed that PLD activity product PA can stimulate pollen tube growth and germination (Potocký *et al.*, 2003; Pleskot *et al.*, 2012). Here, we studied the effect of YFP:NtPLDδ1-5 expressions on the PA distribution in PM using genetically-encoded PA biosensor (mRFP:2xSpo20p-PABD) that we recently described in pollen tubes (Potocký *et al.*, 2014; Sekereš *et al.*, 2017). We cotransformed tobacco pollen with mRFP:2xSpo20p-PABD and individual YFP:NtPLDδs or free YFP and we analyzed the PM fluorescence intensity of PA biosensor in morphologically normal growing pollen tubes by confocal microscopy. Interestingly, only coexpression with YFP:NtPLDδ3 led to significant increase of mRFP:2xSpo20p-PABD signal at PM compared to free YFP control (Fig. 7a). Because transient cotransformation always yields cells expressing two constructs at different levels, making direct fluorescence intensity comparisons challenging, we performed a ratiometric analysis to quantitatively assess the effect of PLDδs expression on PA levels. Here we took advantage of concomitant plasmalemma (PA-specific) and nuclear (PA-nonspecific) localization of mRFP:2xSpo20p-PABD and calculated relative PA levels as a ratio between PM and nuclear fluorescence intensity (Nakanishi *et al.*, 2004; Zeniou-Meyer *et al.*, 2007; Potocký *et al.*, 2014). Expression of YFP:NtPLDδ3 increased relative PA levels at PM 6.9x relative to free YFP (Fig. 7b) and moreover, clear positive correlation exists between the expression level of YFP:NtPLDδ3 and mRFP:2xSpo20p-PABD (Fig. S5a), thus confirming initial microscopic observations.

**Fig. 7.**
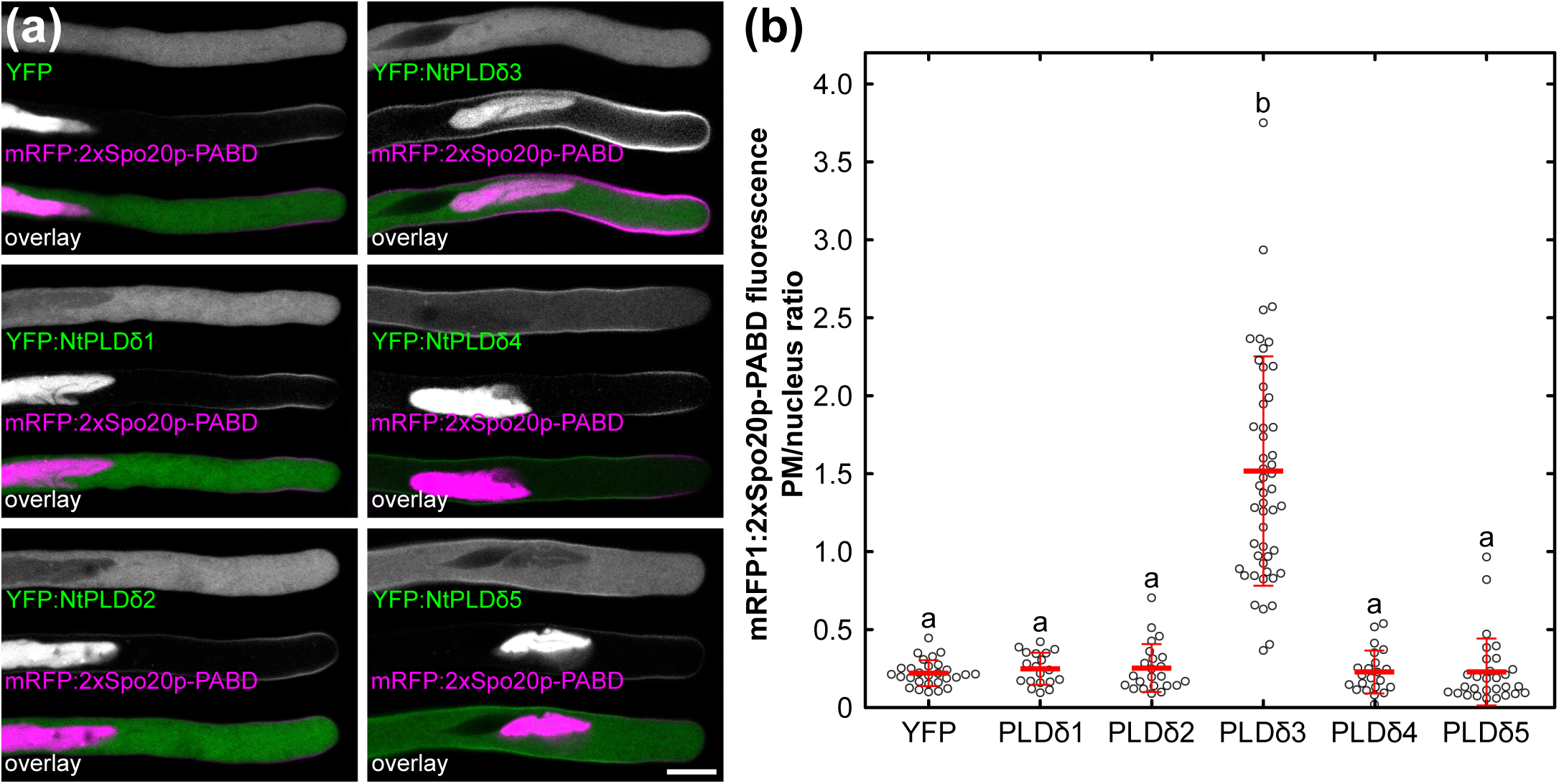
NtPLDδ3 shows high intrinsic PLD activity *in vivo*. **(a)** Colocalization of free YFP and YFP:NtPLDδ1-5 with PA marker mRFP:2xSpo20p-PABD. Bar, 10 μm. **(b)** Quantification of PM PA levels in cells cotransformed with 2.5 µg of free YFP or YFP:NtPLDδ1-5 with 0.5 µg of PA marker mRFP:2xSpo20p-PABD. Pollen grains were analyzed 8 to 10 h after transformation. Data show values for individual cells and mean ± SD (n > 20, three independent experiments). Different letters indicate significant differences between samples (P < 0.001). PLD, phospholipase D.

Having observed that high expression of NtPLDδ3 caused membrane apical invaginations and the level of its expression correlated with PA plasma membrane signal, we tested whether the formation of these abnormal membrane structures is dependent on catalytic activity of NtPLDδ3. We prepared YFP:NtPLDδ3 versions with point mutations in the first (YFP:NtPLDδ3 K351R), second (YFP:NtPLDδ3 K693R) or both (YFP:NtPLDδ3 K351R K693R) HxKxxxxD motives of putative active site by replacing lysine with arginine according to Sung *et al*. (1997) (see also Fig. S3). We cotransformed tobacco pollen with mRFP:2xSpo20p-PABD and individual mutated versions of YFP:NtPLDδ3. Using the ratiometric approach described above, we proved that none of mutated YFP:NtPLDδ3 variants were able to increase PM mRFP:2xSpo20p-PABD signal confirming their catalytic inactivity (Fig. 8a,b and Fig. S5b). Significantly, we detected strong accumulation of mRFP:2xSpo20p-PABD signal in the membrane invaginations upon WT YFP:NtPLDδ3 overexpression, while we never observed either the formation of membrane invaginations or the PA accumulation upon overexpression of inactive YFP:NtPLDδ3 variants (Fig. 8a,b and Fig. S5c).

**Fig. 8.**
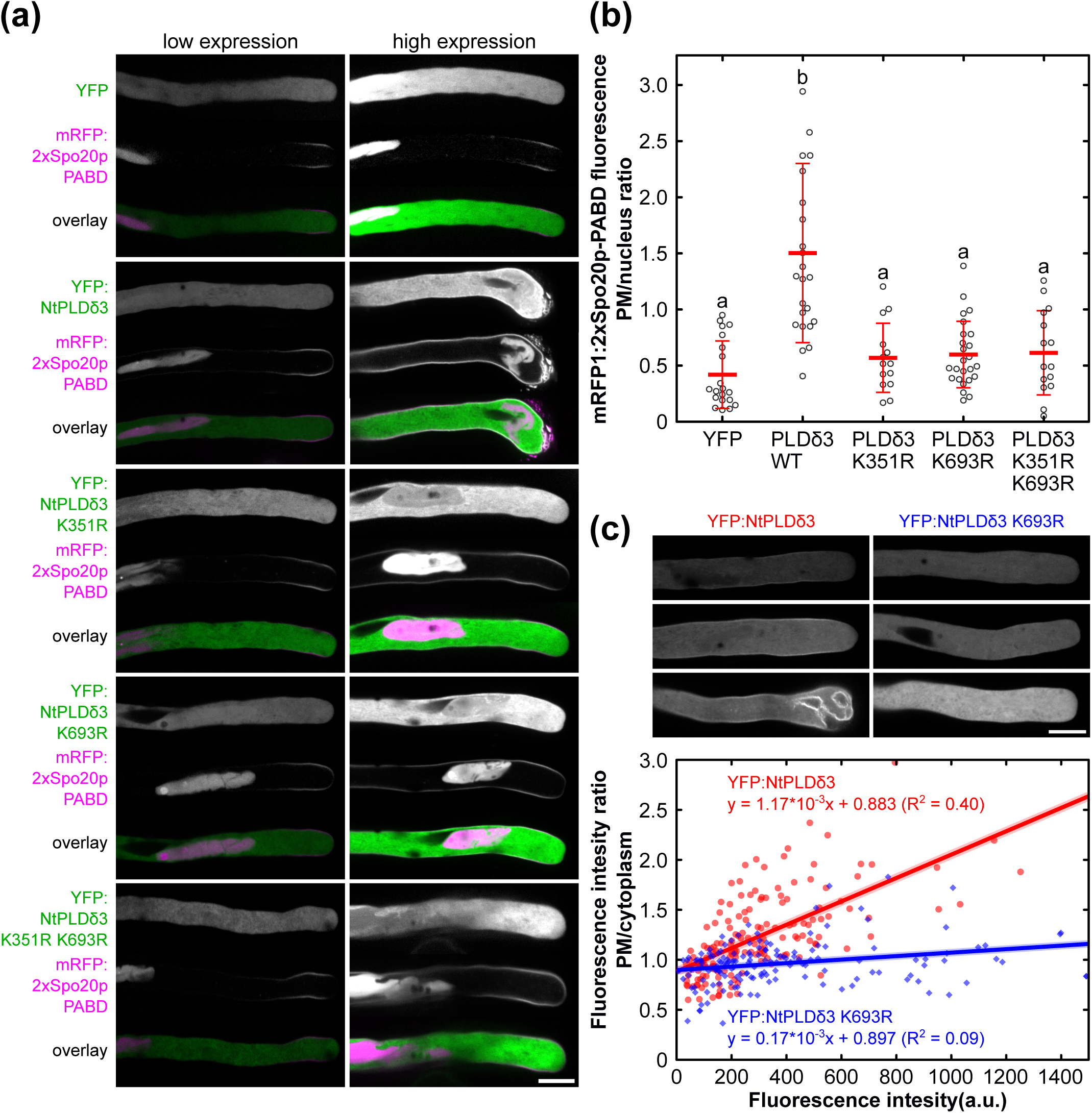
Elevated activity of NtPLDδ3 creates positive feedback enhancing its association with PM and resulting in membrane invaginations. **(a)** *In vivo* PLD activity of WT YFP:NtPLDδ3 and versions with point mutations in putative active site residues. Constructs coding WT and mutated YFP:NtPLDδ3 (2.5 µg) were cotransformed with PA marker mRFP:2xSpo20p-PABD (0.5 µg) and imaged 8 to 10 h after transformation. Typical cells with low and high PLDδ3 levels are shown. **(b)** Quantitative analysis of *in vivo* PLD assay. Data show values for individual cells and mean ± SD (n > 14, two independent experiments). Different letters indicate significant differences between samples (P < 0.001). **(c)** PM binding of NtPLDδ3 is dependent on phospholipase D activity. Upper, typical examples of pollen tubes expressing different levels of WT YFP:NtPLDδ3 and YFP:NtPLDδ3 K693R. See Fig. S6 for more examples. Lower, scatter plot showing positive correlation between PM localization and expression level for WT YFP:NtPLDδ3 but not YFP:NtPLDδ3 K693R. Pollen was transformed with 1-2.5 µg of plasmid DNA and imaged 8 to 10 h after (n > 150, six independent experiments). Bars, 10 μm. PLD, phospholipase D.

Since PA produced by PLD is known to play a role in actin organization in tobacco pollen tubes, we tested the effect of mRFP:NtPLDδ3 expression on actin using tobacco pollen tubes expressing actin marker Lifeact:YFP. However, we did not observe any changes in actin organization upon mRFP:NtPLDδ3 expression compared to mRFP:NtPLDδ3 K693R or Lifeact:YFP only (Fig. S5d) suggesting that NtPLDδ3 is not involved in actin organization. Intriguingly, during the microscopic analyses of wt and inactive NtPLDδ3 variants, we noticed that PM binding of YFP:NtPLDδ3 seems to be positively regulated by phospholipase D catalytic activity, as relative PM NtPLDδ3 signal increases with the expression level for the wt but not inactive variant (Fig. 8c, Fig. S6). Collectively, these experiments showed that PA formed by NtPLDδ3 is required for the proper regulation of pollen tube morphology and positively affects NtPLDδ3 PM binding via a positive feed-back mechanism.

## Discussion

### Evolutionary conserved roles of PLDδ class in male gametophyte

We have provided a genome-wide identification and comparative analysis of the PLD gene family in several *Solanaceae* species representing the first such study in asterids, where systematic PLD analysis has not yet been performed. Our data, documenting 13 C2-PLD and 2 PXPH-PLD genes in diploid *Solanaceae* genome, corroborate the general picture of plant PLD evolution, where plant specific C2-PLD subfamily has significantly expanded during evolution. The mapping of expression data onto the PLD phylogeny showed remarkable evolutionary conservation of expression patterns and suggested that in the broadest terms, 3-exon PLDα class is preferentially expressed in sporophyte, while 10-exon PLDδ class shows preferential expression in pollen. Interestingly, although both PLDα and δ classes are clearly present already in mosses and lycophytes (Pleskot *et al.*, 2012), no apparent isoform specific expression preference between sporophytes and gametofors was found (Potocký, unpublished observations), suggesting that PLDδ isoforms acquired novel functions later, after the dawn of land plants. Evolutionary conserved divergence and expression of isoforms is retained in angiosperms even within the individual classes, i.e. clear trends can be seen for three PLDδ subclasses (corresponding to NtPLDδ1/2, NtPLDδ3 and NtPLDδ4/5). The presence of a single PLDδ gene in Arabidopsis (which belongs to NtPLDδ4/5 subclass) is most probably the outcome of secondary gene loss, as closely-related *Brassica napus* clearly contains members of all three subclasses (Lu *et al.*, 2019). PLDδ gene family composition in Arabidopsis is therefore nonstandard from an evolutionary point of view, strengthening the need for the study of PLDδ function in male gametophyte in other plant model species. In addition to many pharmacological studies showing the importance of PA generated by PLD for pollen tube growth (Potocký *et al.*, 2003; Zonia & Munnik, T., 2004; Monteiro *et al.*, 2005; Pleskot *et al.*, 2010, 2012), the importance of PLDδ family members in pollen tube growth regulation was recently described in pear by Chen *et al*. (2018), who found that the activity of pear PLDδ1 (ortholog of NtPLDδ3) is important for the self-incompatibility signalling pathways.

### Distinct membrane-targeting and activity regulation mechanisms of different PLDδ isoforms

Despite their high degree of overall sequence similarity, members of three PLDδ subclasses displayed different localization patterns and membrane dynamics behaviour. Sequences of tobacco PLDδ proteins do not contain transmembrane helices or typical sites for covalent attachment of lipid-anchors with the exception of NtPLDδ1, where non-canonical myristoylation site is consistently predicted (Potocký, unpublished observation). The membrane-binding mode of many peripheral PM proteins is usually mediated through single domain or sequence motif (Cho & Stahelin, 2005). On the other hand, the membrane attachment of protein with lipid-modifying enzyme activities is often controlled by multiple factors, ensuring precise (and often reversible) control of protein-membrane association. This was shown by Helling *et al*. (2006), who demonstrated that for correct PM localization of tobacco PLC2, the presence of C2 and EF domains is both necessary and sufficient. Similarly, the localization of Arabidopsis PIP5K2 to PM and nucleus is dependent on the linker domain and nuclear localization motif located inside it (Stenzel *et al.*, 2012; Gerth *et al.*, 2017).

Our data suggest that the interaction of various PLDδ members with the PM is even more complex multi-step process involving lipid-binding domains, short sequence motifs, active site residues and putative protein partners. Based on our protein localization and domain deletion/swapping experiments together with *in vivo* PLD activity measurements, we propose three basic scenarios describing the regulation of biological functions for distinct PLDδ subclasses (Fig. 9). In this model, NtPLDδ4/5 isoforms are strongly attached to the PM via the combination of N-terminal phospholipid- and ion-binding C2 domain and short internal motif (VLREGTEI in NtPLDδ4) located within the catalytic domain, that is present only in this subclass. In order to prevent the uncontrollable cleavage of phospholipids, NtPLDδ4/5 isoforms have very low or zero basal PLD activity and the stimulation/inhibition of phospholipase activity would be the major regulatory step. Conversely, NtPLDδ3 possesses high basal activity and its association with PM is much weaker. Notably, this subclass lacks the VLREGTEI motif crucial for PM targeting of NtPLDδ4/5 subclass, instead its PM localization is dependent on phospholipase activity, thus allowing feed forward regulatory loop. It is tempting to speculate that NtPLDδ3 binding to PM is partly due to covalent intermediate with the phosphatidyl moiety connected to the catalytic PLD site as proposed by Dhonukshe *et al*. (2003). Lastly, NtPLDδ1/2 are predominantly cytoplasmic and we hypothesize that for this subclass, both translocation to the PM and enzyme activation will be employed for regulation.

**Fig. 9.**
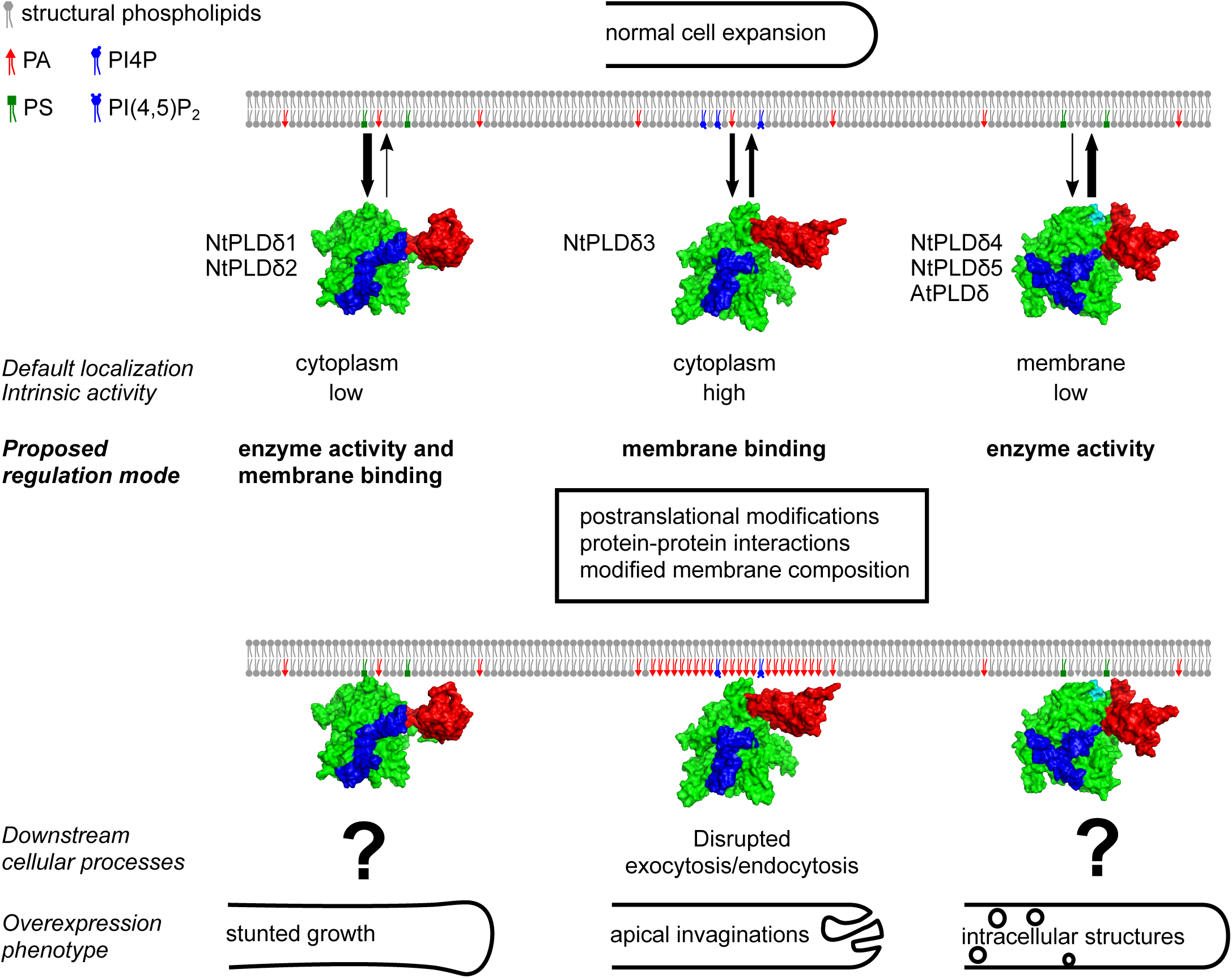
Schematic model describing three scenarios how the biological functions of phospholipases D can be differentially controlled by modulating their innate ability to bind target membranes and/or their intrinsic enzymatic activity. Schematic PLDδ structure shows N-terminal C2 domain in red, central catalytic domain in green and C-terminal domain in blue. See text for more details. PA, phosphatidic acid; PI4P, phosphatidylinositol 4-phosphate; PI4,5P_2_, phosphatidylinositol 4,5-bisphosphate; PLD, phospholipase D; PS, phosphatidylserine.

While the molecular nature of NtPLDδ activity modulators remains elusive, regulation by calcium, phosphoinositides, reactive oxygen species (ROS), phosphorylation and through regulatory proteins are all plausible candidates. Indeed, both calcium and PIP_2_ stimulate Arabidopsis *in vitro* PLDδ activity in dose-dependent manner (Qin *et al.*, 2002) and ROS were implicated in PLDδ regulation *in vivo* for both sporophyte tissues and pollen (Zhang *et al.*, 2003; Potocký *et al.*, 2012). Similarly, phosphorylation was reported to regulate PIP_2_-dependent PLD activity in Brassica (Novotná *et al.*, 2003) and trimeric G-protein α subunit binds and regulates tobacco PLDα (Lein & Saalbach, 2001).

### The roles of PA produced by NtPLDδ3 in pollen tube growth

Comparative analysis of overexpression and knock-down phenotypes together with *in vivo* PLD activity assays clearly identified NtPLDδ3 as the major PLD isoform in tobacco pollen and suggested specific signalling pathways for PA generated by distinct PLDδ subclasses (Fig. 9). While both overexpression and knockdown of NtPLDδ1/2 produced only mild phenotypes (stunted growth without apparent morphology changes), and major outcome of overexpressed NtPLDδ4/5 was intracellular accumulation of large membrane particles of PM origin, elevated levels of NtPLDδ3 activity led to disturbed cell shape characterized by apical PM invaginations. This phenotype is strongly indicative of disturbed membrane trafficking, suggesting a disbalance between rates of exocytosis and endocytosis. In pollen tubes and other tip-growing cells, exocytosis and endocytic retrieval of proteins and membranes from PM are equally important cellular processes that determine and maintain cell polarity (Qin & Yang, 2011; Grebnev *et al.*, 2017). In fact, endocytosis is extremely abundant in pollen tubes, since the fusion of secretory vesicles with the apical PM provided significantly more membrane material than required for PM extension and the maintenance of proper cell architecture requires internalization of the excess membrane (Derksen *et al.*, 1995; Ketelaar *et al.*, 2008). This may account for up to 90% of membranes in secretory vesicles and even when kiss-and-run exocytosis may represent half of the secretory events (Bandmann *et al.*, 2011), ∼40% of membrane material must be endocytosed from the PM.

In pollen tubes and root hairs, PIP_2_ (and to a lesser extent PI4P) were the major phospholipids implicated in exocytosis and endocytosis so far, as elevation PIP_2_ levels were shown to have profound effects on cell polarity and polar tip growth (Heilmann & Ischebeck, 2016). Remarkably, overexpression of several tobacco or Arabidopsis PIP5K isoforms in pollen tubes produces nearly identical membrane invagination phenotype as NtPLDδ3. Invaginations induced by AtPIP5K6 overexpression are probably formed due to excessive clathrin-dependent membrane invagination, supporting the role for PIP_2_ in promoting early stages of clathrin-dependent endocytosis (Zhao *et al.*, 2010). On the other hand, high levels of PIP5K4/5 led to apical pectin accumulation in addition to PM deformations, suggesting excessive secretion (Ischebeck *et al.*, 2008). Despite phenotypic similarities induced by high levels of PIP_2_ and PA, their molecular targets may differ, because their distribution in pollen tube PM only partially overlaps, with PIP_2_ near the apex and PA further back in the subapical zone (Potocký *et al.*, 2014). PA may therefore be involved in other cell morphogenesis-regulating processes besides direct involvement with vesicle traffic machinery. Proper organization and dynamics of cytoskeleton is crucial for membrane traffic in plant cells and PA plays an important role in actin polymerization (Pleskot *et al.*, 2014) and the other way round, microtubule stabilization modified PLD activity (Potocký *et al.*, 2003; Pejchar *et al.*, 2008). Our experiments and previous study (Pleskot *et al.*, 2010) however suggest that NtPLDδ3 is not directly involved in actin regulation. On the other hand, the connection between PLDδ-class isoforms and ROS signalling pathways is well documented. PA may therefore play a role in controlling ROS production via NADPH oxidases, which are crucial for tip growth in pollen tubes and root hairs (Foreman *et al.*, 2003; Potocký *et al.*, 2007; Kaya *et al.*, 2014; Lassig *et al.*, 2014). Notably, the overexpression of ANXUR1/2, receptor like kinases that control pollen NADPH oxidases, phenocopies the effect of elevated NtPLDδ3 activity (Boisson-Dernier *et al.*, 2013).

In summary, we have shown that of PLDδ class have diverse functions in tobacco pollen, which are reflected by their distinct localization and dynamics, membrane targeting and overexpression phenotypes. Our data highlighted the NtPLDδ3 as the predominant pollen PLDδ isoform, and demonstrated that NtPLDδ3-produced PA is crucial for proper control of PM-related membrane traffic at the cell apex. This indicates that in plant cells, the involvement of phospholipids in the regulation of vesicular transport goes beyond phosphoinositides and future studies will be aimed at the identification and characterization of novel players in the secretory machinery that may be regulated by PA.

## Experimental procedures

### Phylogenetic analysis

PLD homologues were identified using BLAST in NCBI, SolGenomics (Mueller *et al.*, 2005) and Phytozome (Goodstein *et al.*, 2012) databases. Exon-intron structures were curated using SoftBerry FGENESH+ algorithm (www.softberry.com) with the aid of experimentally verified sequences. The alignment was constructed using the MAFFT E-INS-I algorithm (Katoh & Standley, 2013) in Jalview (Waterhouse *et al.*, 2009). Gaps and non conserved regions were removed from the alignment, giving the matrix of 85 taxa and 649 positions. Bayesian phylogeny inference was performed using MrBayes (Ronquist *et al.*, 2012) with a WAG + G + I model, where the analysis was performed in four runs with four chains and 1000000 generations, and trees were sampled every 100 generations. All runs approached stationarity after 250000 generations, which were omitted from the final analysis. Maximum likelihood (ML) was calculated using PhyML (Guindon *et al.*, 2010) using LG + G + I model with all parameters estimated from data. ML tree branch values were calculated with approximate likelihood ratio test with SH-like support.

### Gene expression analyses

Total RNA was isolated from tobacco pollen grains and pollen tubes using Qiagen RNAeasy Kit and Turbo DNA-free Kit (Applied Biosystems) was used for DNA removal. cDNA synthesis was carried out using Transcriptor High Fidelity cDNA Synthesis Kit (Roche) with anchored-oligo(DT)18 primer according to manufacturer’s instructions.

Semi-quantitative RT–PCR was done with NtPLDδ1-5 gene-specific oligonucleotides E1-E12 (Supporting Information Table S1) designed to span an intron in the corresponding genomic DNA sequence. Actin7 (Bosch *et al.*, 2005) was used as load control. Amplification conditions were 94 °C for 30□s, 58 °C for 30□s, and 72 °C for 2□min for 35 cycles.

Transcript abundance of tobacco PLD isoforms in pollen was assessed by reanalyzing RNA-seq Illumina data generated by Conze *et al*. (2017) according to Gosh & Chan (2016). Briefly, raw data reads were analyzed using fastQC and mapped to a reference gene-set (TN90 cultivar) using tophat. Cufflinks was used to assemble and reconstruct the transcriptome and calculate transcript per million (TPM) expression values. Analysis of the PLD gene expression in pollen, leaves and roots of Arabidopsis, poplar, tomato and rice was performed using the CoNekT database (Proost & Mutwil, 2018). Additional data for tobacco, poplar and wine were obtained using GEO tool (Barrett *et al.*, 2013) and from Zhao *et al*. (2016).

### Antisense ODN design

NtPLDδ1-5 coding sequences were analyzed using Soligo software 5 for suitable antisense ODNs sites according to Potocký *et al*. (2019). Two best-scoring antisense ODNs and corresponding sense control ODNs were synthesized and tested for their effect on tobacco pollen tube growth, and the more effective pair was used for further experiments.

### Molecular cloning

All coding sequences were amplified flanked by NgoMIV and ApaI sites unless otherwise stated by PCR with Phusion DNA polymerase (NEB).

#### YFP:PLDδ/PLDδ:YFP/mRFP:PLDδ

The NtPLDδ1-5 and AtPLDδ coding sequences were generated from *Nicotiana tabacum* pollen tube and *Arabidopsis thaliana* Col-0 flower cDNA, respectively using the specific primers PP1-PP18 (Supporting Information Table S1). To create NtPLDδ3 K351R and NtPLDδ3 K693R, megaprimer MP351-F was generated first from YFP:NtPLDδ3 as a template using PP7/PP19 primers and used in the second step as forward primer together with reverse primer PP8 (for NtPLDδ3 K351R) and megaprimer MP693-R using PP20/PP8 primers and used in the second step as reverse primer together with forward primer PP7 (for NtPLDδ3 K693R). To generate NtPLDδ3 K351R K693R, megaprimers MP351-F/MP693-R were used.

#### YFP:NtPLDδ4 domains

NtPLDδ4-N, NtPLDδ4-ΔN, NtPLDδ4-C, NtPLDδ4-ΔC, NtPLDδ4-ΔNΔC domains were generated from YFP:NtPLDδ4 as a template using primers PP10, PP11 and PP21-PP24. To create NtPLDδ4-NC, NtPLDδ4-ΔSHWHD and NtPLDδ4-ΔVL-EI, megaprimer was generated first from YFP:NtPLDδ4 as a template using forward primers PP25, PP26 and PP27, respectively and reverse primer PP11 and used in the second step as reverse primer together with forward primer PP10.

#### YFP:NtPLDδ2/NtPLDδ4 chimeric proteins

First, set of four megaprimers (MP) was generated using primers PP28-PP31. MPδ24x-F (using PP3/PP28) and MPδx42-R (using PP29/PP4) were amplified from YFP:NtPLDδ2 as a template while MPδ42x-F (using PP10/PP30) and MPδx24-R (using PP31/PP11) were amplified from YFP:NtPLDδ4. Then NtPLDδ242 (using MPδ24x-F/MPδx42-R), NtPLDδ244 (using MPδ24x-F/PP11) and NtPLDδ442 (using/and MPδx42-R) were generated from YFP:NtPLDδ4 as a template. NtPLDδ424 (using MPδ42x-F/MPδx24-R), NtPLDδ422 (using MPδ42x-F/PP4) and NtPLDδ224 (using PP3/MPδx24-R) were generated from YFP:NtPLDδ2 as a template.

Amplified products were introduced into the multiple cloning sites of the pollen expression vector pWEN240, pHD32 or pHD222 for N-terminal and C-terminal YFP and N-terminal mRFP fusion, respectively (Klahre *et al.*, 2006).

#### HA-NtPLDδ2-4

First, NtPLDδ2-4 were amplified using primers PP32-PP37 carrying overhang for PP38 and NotI site from YFP:NtPLDδ2-4 as templates. In a second step, forward primer with SalI site, Kozak sequence and HA-tag PP38 and reverse primers PP33, PP35 and PP37 and NtPLDδ2-4 products from the first step were used as templates for amplification. Final products were introduced into vector pTNT (Promega) via SalI and NotI sites.

#### mRFP:2xSpo20-PABD

Spo20-PABD coding sequence was amplified from YFP:Spo20-PABD (Potocký *et al.*, 2014) as a template using primers PP39 and PP40 carrying SpeI/XmaI sites. Amplified product was introduced into SpeI/XmaI sites of mRFP:Spo20-PABD resulting in creation of mRFP:2xSpo20-PABD.

### Pollen transformation and the microscopic analysis of pollen tube growth

Tobacco pollen grains germinating on solid medium were transformed by particle bombardment using a helium-driven particle delivery system (PDS-1000/He; Bio-Rad) as described previously (Kost *et al.*, 1998). Particles were coated with 0.5-5 μg DNA. For live-cell imaging, pollen tubes were observed with a spinning disk confocal microscope (Yokogawa CSU-X1 on Nikon Ti-E platform) equipped with a 60X Plan Apochromat objective (WI, NA = 1.2) and Andor Zyla sCMOS camera. 488 nm laser excitation together with 542-027 nm single band filter (Semrock Brightline) were used for fluorescence collection of YFP. Laser and camera settings were kept constant, allowing for comparative imaging.

FRAP analyses were performed using Zeiss LSM880 equipped with C-Apochromat 40x/1.2WI objective. YFP fluorescence was excited by a 514 nm laser and the emission at 520–590 nm was recorded using a GaAsP detector. Raw images were corrected for pollen tube growth, and intensities were normalized as follows: It = [(At / Ct) - (Ab / Cb)] / [Av - (Ab / Cb)], where At and Ct are intensities in the bleached and control areas, respectively, at time t; Ab and Cb are the intensities in the bleached and control areas, respectively, immediately after the bleaching; and Av is the average At/Ct intensity ratio in the five frames before bleaching. FRAP curves were fitted and analyzed using custom R script with the single and double exponential equations F_single_(x) = A * (1 - exp-B*x) or F_double_(x) = A1 * (1 - exp-B1*x) + A2 * (1 - exp-B2*x).

Figures were processed with ImageJ (rsbweb.nih.gov/ij/) and GIMP (www.gimp.org) and assembled in Inkscape (www.inkscape.org).

### Homology modelling

NtPLDδ2-4 homology models were produced by Robetta (Kim *et al.*, 2004), evaluated with Prosa (prosa.services.came.sbg.ac.at/prosa.php), WhatIf (swift.cmbi.ru.nl/servers/html/index.html) and PSVS (psvs.nesg.org/) servers. Electrostatic potentials were calculated using APBS (Baker *et al.*, 2001). Blue, red and white color represents positive, negative and neutral potential, respectively. Pymol (www.pymol.org/) was used to visualize the structures.

### Protein–lipid overlay assay

The protein–lipid overlay assay was performed using N-terminally HA-tagged NtPLDδ2-4 and lipid strips (Echelon Biosciences). The constructs were used as a template for *in vitro* coupled transcription/translation reactions using the TNT® SP6 High-Yield Wheat Germ Protein Expression System (Promega) in a total volume of 50 μl and the reaction product was used for the protein–lipid overlay assay as described previously in Kubátová *et al*. (2019).

## Acknowledgments

This work was supported by the Czech Science Foundation grants GA17-27477S to P.P. and GA19-21758S to M.P. Part of income of V.Ž. is supported by the Ministry of Education, Youth and Sports of CR from European Regional Development Fund-Project “Centre for Experimental Plant Biology”: No. CZ.02.1.01/0.0/0.0/16_019/0000738. IEB Imaging Facility is supported by Operační program Praha – Konkurenceschopnost Moderní přístroje pro výzkum rostlin CZ.2.16/3.1.00/21519 and by the project of Ministry of Education, Youth and Sports “National Infrastructure for Biological and Medical Imaging (Czech-BioImaging – LM2015062).

The authors thank Prof. Benedikt Kost (University of Erlangen-Nuremberg, Erlangen, Germany) who kindly provided us constructs for the transient expression in tobacco pollen. The authors also thank the developers of the Linux operating system and the open-source software used in preparation of this study, particularly ImageJ, Inkscape, Gimp and Gnumeric.

## Supporting Information

**Fig. S1** Colocalization of tobacco YFP:NtPLDδ1-5 with FM4-64 and localization of NtPLDδ1-5 and AtPLDδ tagged with YFP on C-terminus.

**Fig. S2** Comparison of single exponential vs double exponential fit of FRAP data.

**Fig. S3** Multiple alignment of NtPLDδ1-5 and AtPLDδ.

**Fig. S4** Membrane-binding properties of NtPLDδ1-5 in cells with different expression levels.

**Fig. S5** Changes in pollen tube morphology caused by NtPLDδ3 overexpression are caused by elevated phospholipase D activity and are independent on actin cytoskeleton.

**Fig. S6** Catalytically inactive NtPLDδ3 cannot efficiently bind PM and induce membrane invaginations.

**Table S1** List of primers used in this study.

## Author contribution

P.P. and M.P. conceived the study, designed and performed all molecular biology, bioinformatic and microscopic experiments, analyzed the data and wrote the manuscript. J.S. performed the lipid-binding analyses and participated in writing of the manuscript. O.N. took part in FRAP analyses. V.Ž. participated in writing of the manuscript. All authors approved the final version of the manuscript.

