## Supporting Information for "Functional analysis of phospholipase Dδ family in tobacco pollen tubes"

Article title: Functional analysis of phospholipase Dδ family in tobacco pollen tubes

The following Supporting Information is available for this article:

**Fig. S1**	Colocalization of tobacco YFP:NtPLDδ1-5 with FM4-64 and localization of NtPLDδ1-5 and AtPLDδ tagged with YFP on C-terminus.

**Fig. S2**	Comparison of single exponential vs double exponential fit of FRAP data.

**Fig. S3**	Multiple alignment of NtPLDδ1-5 and AtPLDδ.

**Fig. S4**	Membrane-binding properties of NtPLDδ1-5 in cells with different expression levels.

**Fig. S5**	Changes in pollen tube morphology caused by NtPLDδ3 overexpression are caused by elevated phospholipase D activity and are independent on actin cytoskeleton.

###### Fig. S6 Catalytically inactive NtPLDδ3 cannot efficiently bind PM and induce membrane invaginations.

**Table S1** List of primers used in this study.

###### Fig. S1 Colocalization of tobacco YFP:NtPLDδ1-5 with FM4-64 and localization of NtPLDδ1-5 and AtPLDδ tagged with YFP on C-terminus. (a) Colocalization of YFP-tagged NtPLDδ1-5 (green) and endocytic dye FM4-64 (magenta) in tobacco pollen tubes. (b) Localization of NtPLDδ1-5 and AtPLDδ tagged with YFP at C-terminus in actively growing pollen tubes. Pollen was transiently transformed with 1 μg of DNA and analyzed 8 to 10 h after transformation. Bars, 10 μm. PLD, phospholipase D.

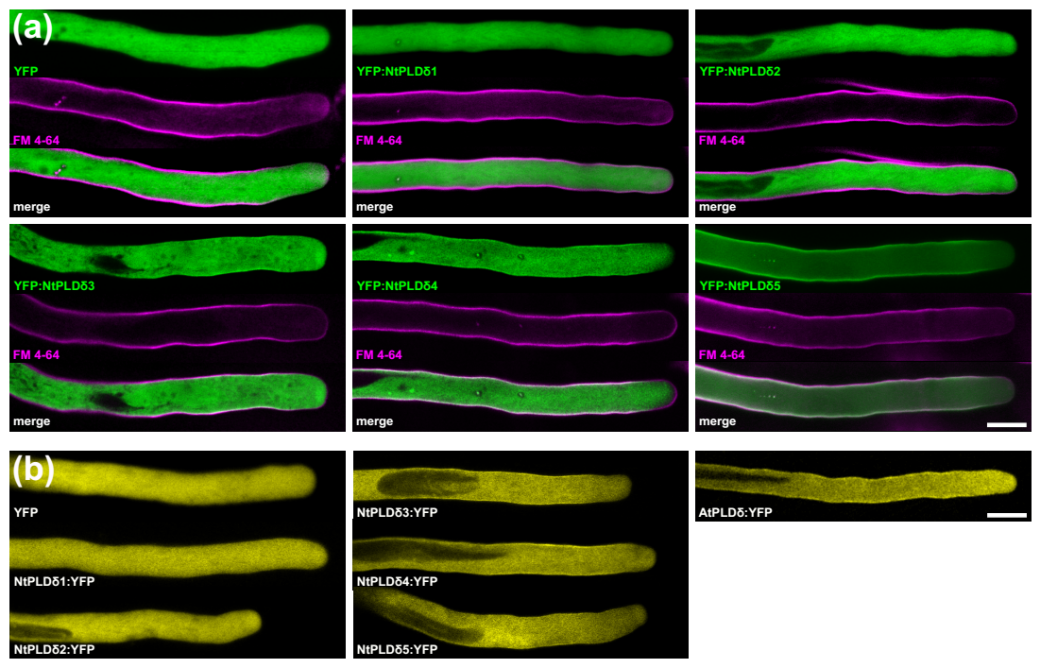

###### Fig. S2 Comparison of single exponential vs double exponential fit of FRAP data. Experimental data (shown as mean ± SEM) were fitted with either single exponential (Intensity = Mf*(1-e^-K*time^), blue line) or double exponential (Intensity = F*(1-e^-Kf*time^) + S*(1-e^-Ks*time^), red line) equation. Final fit (boxed) was chosen based on comparisons of residuals plots, RMSE values, the capacity of individual (fast/slow) parameters to explain the majority of experimental data and standard errors of fitted parameters. Half-times were computed using equation T_1/2_ = ln(2)/K, mobile fraction Mf was obtained using the equation Mf = F + S for double exponential fit. PLD, phospholipase D.

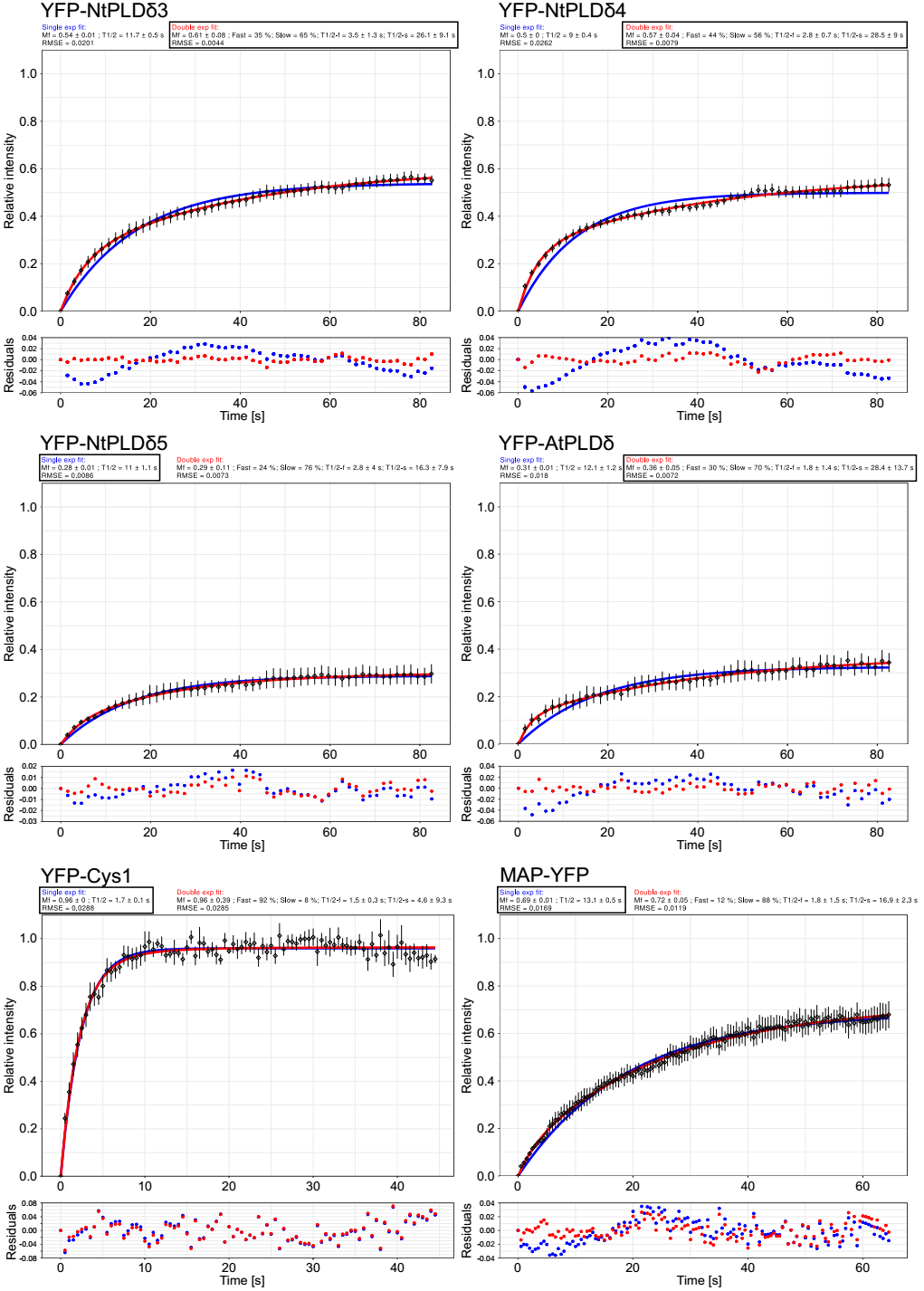

###### Fig. S3 Multiple alignment of NtPLDδ1-5 and AtPLDδ. The presented data are an excerpt from a larger source alignment that served as input for the phylogenetic analysis shown in Fig. 1. Boundaries between N-terminal C2 domain, central catalytic domain and C-terminal domain are indicated by red lines with triangles. Two sequence insertions specific for membrane-bound PLDs are boxed and active site lysine residues used for generation of catalytically-inactive mutants are marked with asterisks. At, *Arabidopsis thaliana*; Nt, *Nicotiana tabacum*; PLD, phospholipase D.

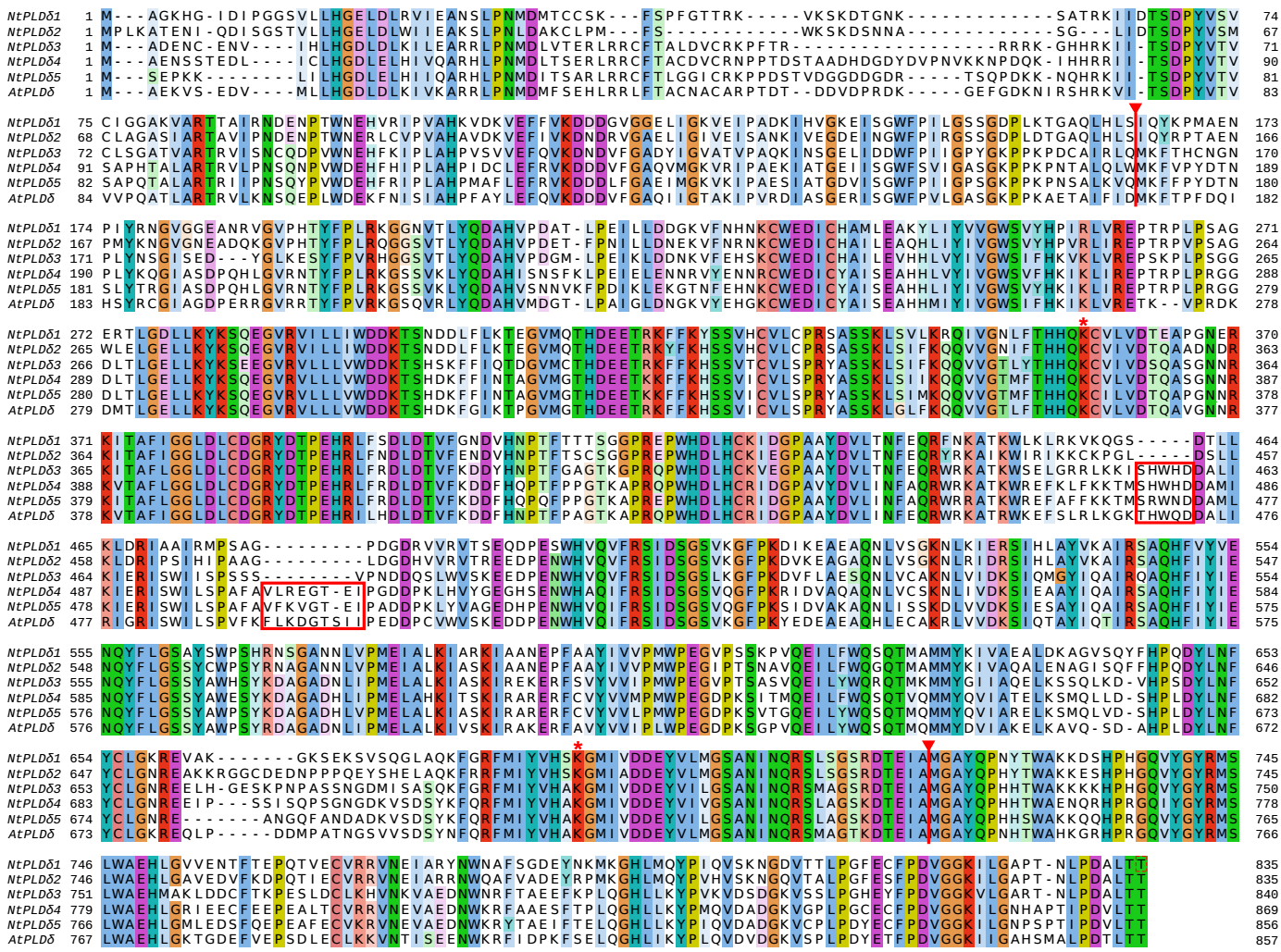

###### Fig. S4 Membrane-binding properties of NtPLDδ1-5 in cells with different expression levels. (a) Typical localization and phenotypes of pollen tubes (PT) expressing various levels of free YFP and YFP-tagged NtPLDδ1-5 (cells correspond to those shown in Fig. 6a). Bar, 10 μm. (b) Relative fluorescence intensity across the PT tip (orange lines), across the PT subapex (purple lines) or across the PT shank (green lines) for pollen tubes shown in Fig. 6a. Fluorescence was measured using the line scan tool in ImageJ across 14 μm-long regions indicated in (a) with line width set to 5 pixels. Pollen was transformed with 0.5 (for low expression level) or 5 μg (for high expression levels) of DNA and analyzed 8 to 10 h after transformation. PLD, phospholipase D.

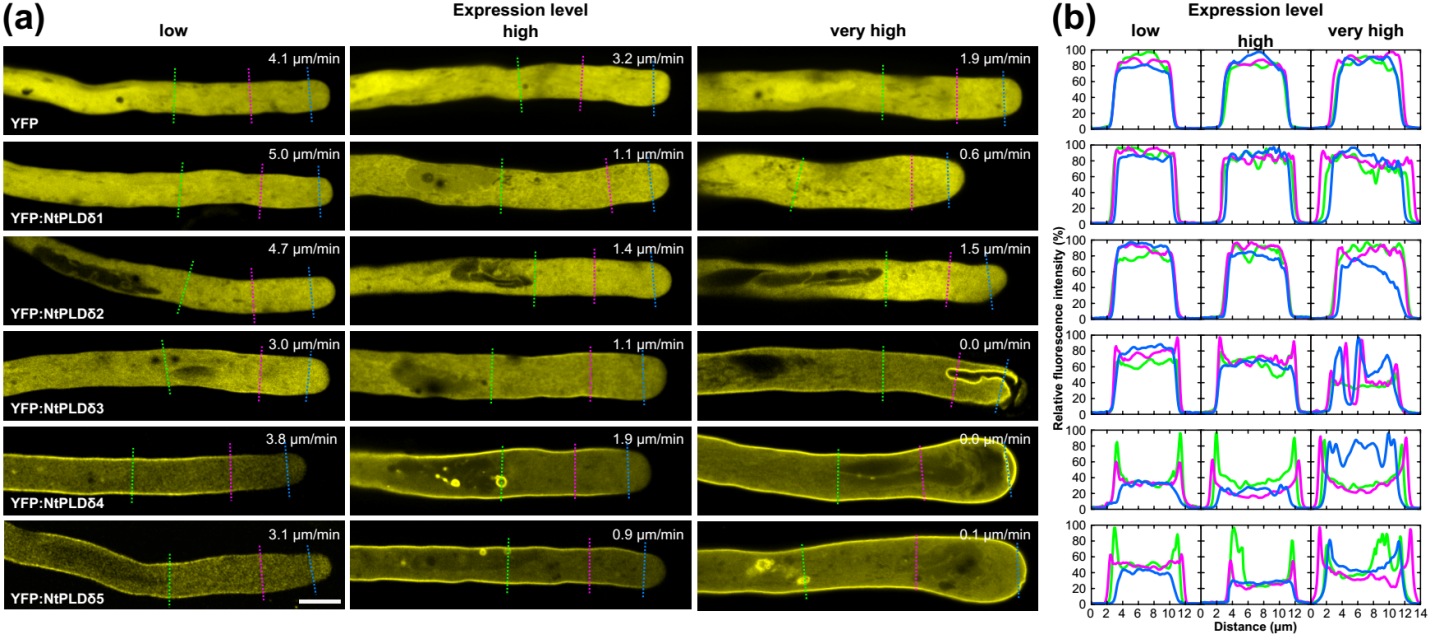

###### Fig. S5 Changes in pollen tube morphology caused by NtPLDδ3 overexpression are caused by elevated phospholipase D activity and are independent on actin cytoskeleton. (a,b) Analysis of *in vivo* PLD activity in cells expressing (a) free YFP or YFP:NtPLDδ1-5, (b) YFP:NtPLDδ3 with mutations in putative catalytic site residues, see also Fig. S3 for details. Pollen was co-transformed with 0.5 µg of mRFP1:2xSpo20p-PABD (read-out for in PA) together with 2.5 µg of free YFP or various YFP:NtPLDδ constructs and imaged 8 to 10 h after. Scatter plots show the experimental data for individual pollen tube, linear regression with 95% confidence intervals, regression parameters and R^2^ coefficients. The R^2^ value indicates the strength of a linear relationship between the relative PM levels of PA marker and expression of free YFP or YFP:NtPLDδ constructs. (c) Examples of cells from (a,b) sorted according to increasing expression level. (d) Elevated PA levels caused by the expression of NtPLDδ3 do not change the structure of actin cytoskeleton. 1 µg of WT mRFP:NtPLDδ3 or inactive mutant mRFP:NtPLDδ3 K693R was transformed into tobacco pollen stably expressing actin marker Lifeact:YFP and observed after 8 h. Bars, 10 μm. PLD, phospholipase D.

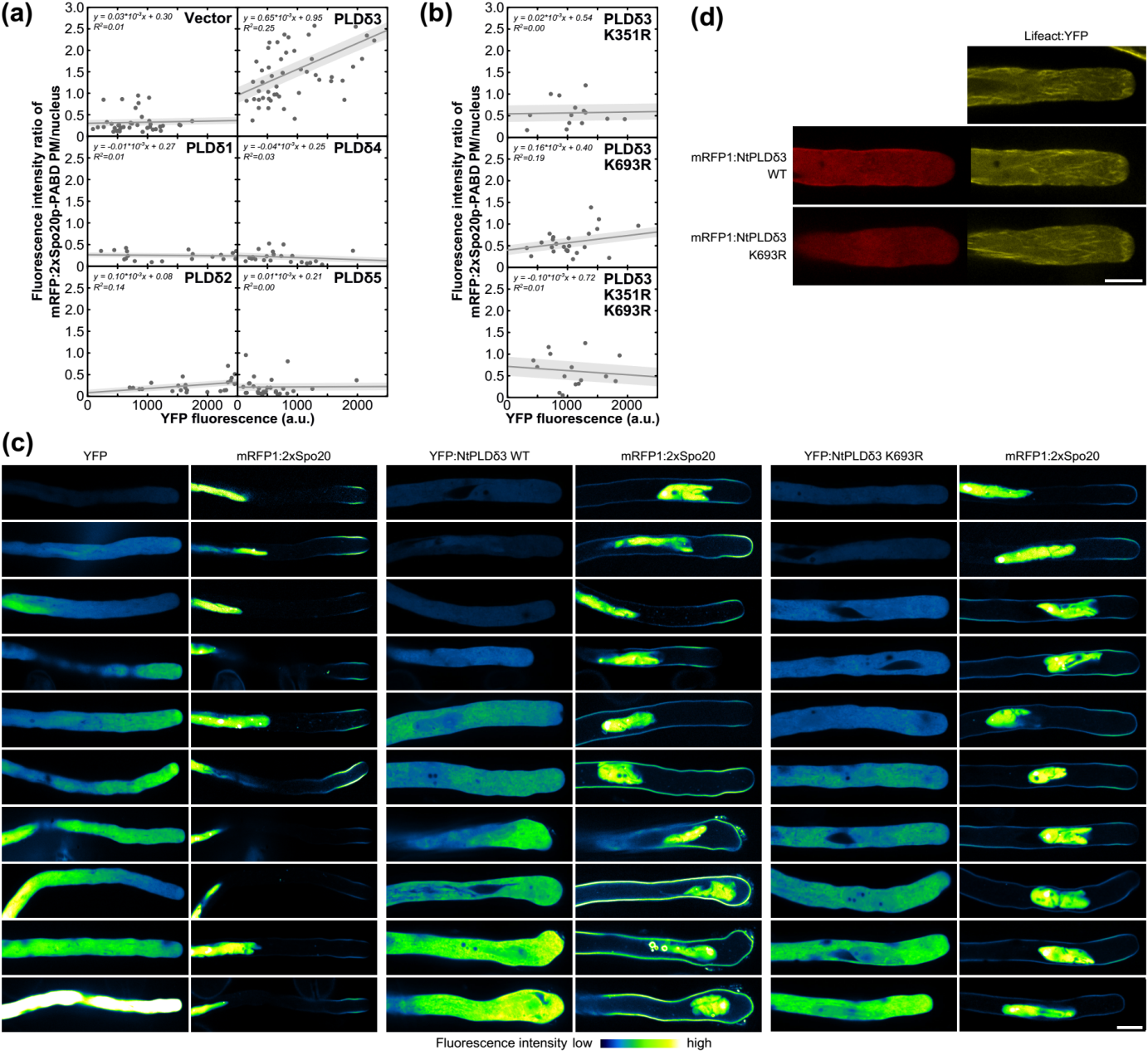

###### Fig. S6 Catalytically inactive NtPLDδ3 cannot efficiently bind PM and induce membrane invaginations. Pollen tube micrographs of cells expressing various levels of YFP:NtPLDδ3 or YFP:NtPLDδ3 K693R imaged with the same acquisition settings. Pollen was transformed with 1-2.5 µg of plasmid DNA and imaged 8 to 10 h after. Examples of cells were sorted according to increasing expression level. Bar, 10 μm. PLD, phospholipase D.

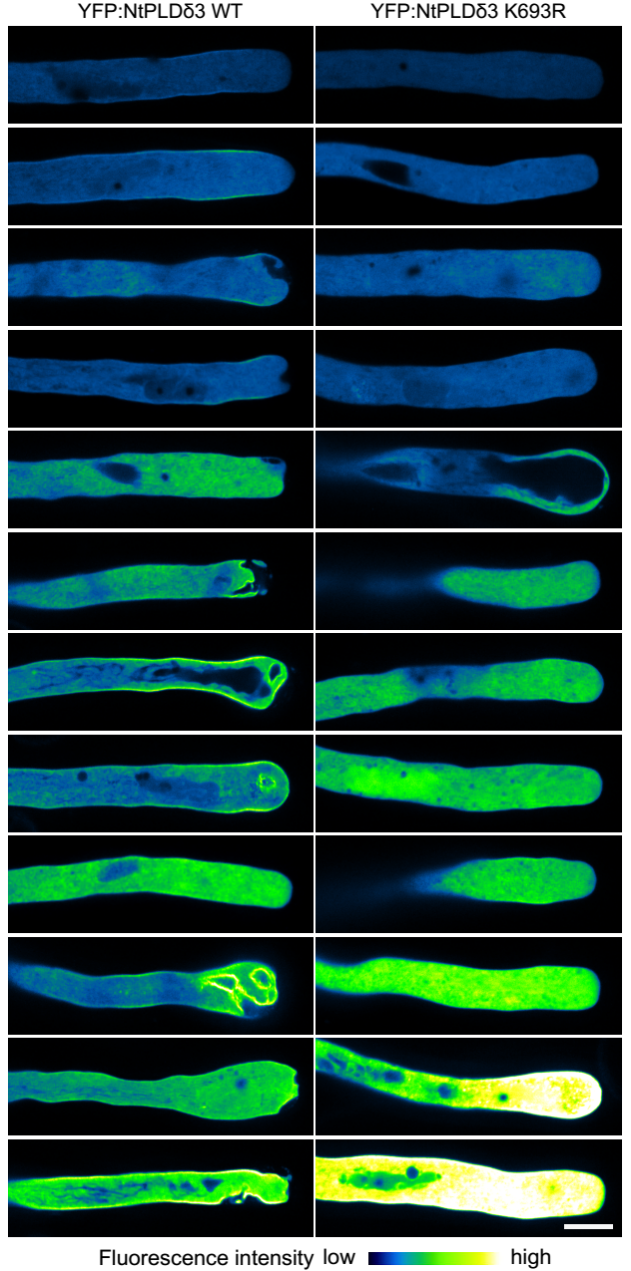

**Table S1**

###### **Supporting Information Table S1.** List of primers used in this study.

| **Label** | **Name** | **Sequence (5**ʼ**- 3**ʼ**)** | **description** |
| --- | --- | --- | --- |
| E1 | NtPLDδ1-F-RT | GGCCTGGATTTATGTGATGG | RT-PCR |
| E2 | NtPLDδ1-R-RT | TTGGACTGGTTTACTGCTTGG | RT-PCR |
| E3 | NtPLDδ2-F-RT | GGTCCTGCTGCTTACGATG | RT-PCR |
| E4 | NtPLDδ2-R-RT | CACGCTTCTTTGCTTCTCTG | RT-PCR |
| E5 | NtPLDδ3-F-RT | TTGAGCAGAGGTGGAGAAAAG | RT-PCR |
| E6 | NtPLDδ3-R-RT | TGAGGAAGCAGGGTTTGG | RT-PCR |
| E7 | NtPLDδ4-F-RT | AGGAAAGCGACAAAATGGAG | RT-PCR |
| E8 | NtPLDδ4-R-RT | TCACCATTACCAGAAGATTGAGAG | RT-PCR |
| E9 | NtPLDδ5-F-RT | AGCGATGGAGGAGAGCAAC | RT-PCR |
| E10 | NtPLDδ5-R-RT | CCATTTGCTTCACGATTTCC | RT-PCR |
| E11 | NtACT7-F-RT | TGCCTATGTTGGTGATGAAGC | RT-PCR |
| E12 | NtACT7-R-RT | ACCATCACCAGAGTCCAACAC | RT-PCR |
| PP1 | NtPLDδ1-F_NgoMIV | ATAGCCGGCGGAACAATGGCTGGAAAACATGG | N-, C-term YFP fusion |
| PP2 | NtPLDδ1-R-S_ApaI | TATGGGCCCGAGTCGTTTAAGTGGTAAGAGCATCAGGA | N-term YFP fusion |
| PP3 | NtPLDδ1-R-NS_ApaI | TATGGGCCCAGTGGTAAGAGCATCAGGA | C-term YFP fusion |
| PP4 | NtPLDδ2-F_NgoMIV | ATAGCCGGCGGAACAATGCCCTTAAAAGCTACTGAGA | N-, C-term YFP fusion |
| PP5 | NtPLDδ2-R-S_ApaI | TATGGGCCCGAGTCGTTTACGTGGTAAGAGCGTCG | N-term YFP fusion |
| PP6 | NtPLDδ2-R-NS_ApaI | TATGGGCCCCGTGGTAAGAGCGTCG | C-term YFP fusion |
| PP7 | NtPLDδ3-F_NgoMIV | ATAGCCGGCGGAACAATGGCGGATGAGAATTG | N-, C-term YFP/RFP fusion |
| PP8 | NtPLDδ3-R-S_ApaI | TATGGGCCCTCATGTGGTCAAAGCATCA | N-term YFP/RFP fusion |
| PP9 | NtPLDδ3-R-NS_ApaI | TATGGGCCCTGTGGTCAAAGCATCAGG | C-term YFP fusion |
| PP10 | NtPLDδ4-F_NgoMIV | ATAGCCGGCGGAACAATGGCGGAAAATTCATCTAC | N-, C-term YFP fusion |
| PP11 | NtPLDδ4-R-S_ApaI | TATGGGCCCGAGTCGTTTATGTTGTCAAAACATCTGGG | N-term YFP fusion |
| PP12 | NtPLDδ4-R-NS_ApaI | TATGGGCCCTGTTGTCAAAACATCTGGG | C-term YFP fusion |
| PP13 | NtPLDδ5-F_NgoMIV | ATAGCCGGCGGAACAATGTCCGAACCTAAAAAGCTC | N-, C-term YFP fusion |
| PP14 | NtPLDδ5-R-S_ApaI | TATGGGCCCGAGTCGTTCAGGTGGTCAACACATCTG | N-term YFP fusion |
| PP15 | NtPLDδ5-R-NS_ApaI | TATGGGCCCGGTGGTCAACACATCTG | C-term YFP fusion |
| PP16 | AtPLDδ-F_XmaI | ATACCCGGGGGAACAATGGCGGAGAAAGTATCG | N-, C-term YFP fusion |
| PP17 | AtPLDδ-R-S_ApaI | ATAGGGCCCTTACGTGGTTAAAGTGTCAGGA | N-term YFP fusion |
| PP18 | AtPLDδ-R-NS_ApaI | ATAGGGCCCCGTGGTTAAAGTGTCAGGAAG | C-term YFP fusion |
| PP19 | NtPLDδ3-K351R-F | TGTGAGTCCACAATCACACACCTCTGATGGTGCGTATAAAGGG | Megaprimer for NtPLDδ3-K351R |
| PP20 | NtPLDδ3-K693R-R | TTTATGATTTATGTACACGCCAGGGGGATGATAGTGGACG | Megaprimer for NtPLDδ3-K351R |
| PP21 | NtPLDδ4-N-R-S_ApaI | ATAGGGCCCTCACCATAGCTGAAGAGCCG | N-term YFP fusion |
| PP22 | NtPLDδ4-ΔN-F_NgoMIV | ATAGCCGGCGGAACAATGAAATTCGTTCCATATGATACA | N-term YFP fusion |
| PP23 | NtPLDδ4-C-F_NgoMIV | ATAGCCGGCGGAACAATGGGTGCCTACCAGCC | N-term YFP fusion |
| PP24 | NtPLDδ4-ΔC-R-S_ApaI | ATAGGGCCCTCAAGCTATCTCTGTGTCCTTTG | N-term YFP fusion |
| PP25 | NtPLDδ4-NC-F | CGGCTCTTCAGCTATGGATGGGTGCCTACCAGCC | Megaprimer for NtPLDδ4-NC |
| PP26 | NtPLDδ4-ΔSHWHD-F | TTCAAGTTGTTTAAGAAAACAATGGATGCTATGATAAAAATTGAGC | Megaprimer for NtPLDδ4-ΔSHWHD |
| PP27 | NtPLDδ4-ΔVL-EI-F | CTAAGCCCTGCTTTTGCTCCAGGGGATGATCCCAAA | Megaprimer for NtPLDδ4-ΔVL-EI |
| PP28 | d2-157_d4-180_MPF24x | TGTATCATATGGAACGAATTTCATCGACAGATGCAGTTGAGC | Megaprimer for NtPLDδ24x |
| PP29 | d4-750_d2-718_MPRx42 | AAGGACACAGAGATAGCTATGGGAGCTTATCAACC | Megaprimer for NtPLDδx42 |
| PP30 | d4-179_d2-158_MPF42x | GCTGTCGGCCTATATTGGATCCATAGCTGAAGAGCCGT | Megaprimer for NtPLDδ42x |
| PP31 | d2-719_d4-751_MPRx24 | AAGGGATACAGAAATAGCTATGGGTGCCTACCAG | Megaprimer for NtPLDδx24 |
| PP32 | NtPLDδ2-F_HA-oh | GATGTTCCAGATTACGCTATGCCCTTAAAAGCTACTGAGAAC | N-term HA fusion |
| PP33 | NtPLDδ2-R-S_NotI | ATAGCGGCCGCTTACGTGGTAAGAGCGTCGGG | N-term HA fusion |
| PP34 | NtPLDδ3-F_HA-oh | TTCCAGATTACGCTATGGCGGATGAGAATT | N-term HA fusion |
| PP35 | NtPLDδ3-R-S_NotI | TAGCGGCCGCTCATGTGGTCAAAGCATCA | N-term HA fusion |
| PP36 | NtPLDδ4-F_HA-oh | TTCCAGATTACGCTATGGCGGAAAATTCATCTAC | N-term HA fusion |
| PP37 | NtPLDδ4-R-S_NotI | TAGCGGCCGCTTATGTTGTCAAAACATCTGGG | N-term HA fusion |
| PP38 | pTNT-HA-overhang_SalI | AAGTCGACGCCGCCACCATGTACCCATACGATGTTCCAGATTACGCTATG | N-term HA fusion |
| PP39 | GA5-Spo20-F_SpeI | GCGACTAGTGGTGCTGGTGCTGGTGCTGGTGCTGGTGCCGGCATG | mRFP:2xSpo20-PABD |
| PP40 | Spo20-R-S_XmaI | TATCCCGGGTCATCTTGTCTTAGTGGCGTCATCG | mRFP:2xSpo20-PABD |
